# Aquaporin-3 amplifies arsenite genotoxicity as a dose-dependent gene-by-environment interaction in human cells

**DOI:** 10.64898/2026.09.23.752800

**Authors:** Shiwei Yin, Bingru Feng, Maryam Vaziripour, Shannon E. Slewitzke, Gail F. Fernandes, Jihye Yun, Christopher I. Amos, Jun Xia

**Affiliations:** Institute of Biosciences and Technology, Texas A&M Health Science Center, Houston, TX 77030, USA; Department of Translational Medical Sciences, Texas A&M University, Houston, TX 77030, USA; Center for Genomic and Precision Medicine, Institute of Biosciences and Technology, Texas A&M Health Science Center, Houston, TX 77030, USA; Center for Epigenetics and Disease Prevention, Institute of Biosciences and Technology, Texas A&M Health Science Center, Houston, TX 77030, USA; Department of Molecular and Human Genetics, Baylor College of Medicine, Houston, TX 77030, USA; Institute for Clinical and Translational Research, Baylor College of Medicine, Houston, TX 77030, USA; Department of Genetics, The University of Texas MD Anderson Cancer Center, Houston, TX 77030, USA; UNM Comprehensive Cancer Center, University of New Mexico, Albuquerque, NM 87131, USA

## Abstract

Chronic exposure to inorganic arsenic affects more than 200 million people, and disease outcome varies widely among individuals with comparable intake. Explanations have centered on arsenic metabolism, which acts on arsenite that is already inside the cell. Uptake sits upstream of metabolism and sets how much arsenite reaches the genome, yet whether it acts as a gene-by-environment modifier of genotoxicity has been asserted far more often than it has been measured. We expressed human aquaporin-3 (AQP3), its point mutants and the paralogues AQP7, AQP9 and AQP10 in three human cell lines, and measured DNA damage as γH2AX by flow cytometry against a damage threshold re-derived from mock-transfected cells in every experiment. In lung fibroblasts, AQP3 raised the γH2AX-high fraction 2.36-fold at 5 μM arsenite. Because a gene-by-environment interaction widens the genotype gap with dose while an independent genotoxin would not, the combination exceeds what the two factors would produce if they simply added by 6.36 percentage points, which is an interaction. The response required the AQP3 sequence, was suppressed by N-acetylcysteine in proportion to arsenite dose, was absent in RKO cells, and was shared with AQP9 and AQP10, which created an arsenite dose-response in a cell line that had none of its own. Low-input RNA sequencing of 135 ten-cell pools located the effect inside the cell: at an identical 1 μM applied dose and within a single batch, AQP3 raised *HMOX1* (induced when trivalent arsenic modifies KEAP1 thiols, and so a report of intracellular rather than applied exposure) 3.3-fold, a shift equivalent to 40% of the 1-to-10 μM interval on the measured dose-response, while AQP3 without arsenite was indistinguishable from untreated. Duplex sequencing detected no consistent change in somatic mutation frequency from arsenite alone at any dose to 10 μM, in an assay that resolved a 26-fold response to N-ethyl-N-nitrosourea; the arsenite-induced mutation frequency at the *EGFR* locus nonetheless differed 1.61-fold by AQP3 genotype. The interaction is therefore not unique to AQP3, and aquaglyceroporin complement belongs alongside metabolism among the host factors that set arsenic susceptibility.

## INTRODUCTION

Inorganic arsenic (iAs) is among the most widespread environmental carcinogens and is classified by the International Agency for Research on Cancer as carcinogenic to humans (Group 1) (1, 2). Chronic exposure through contaminated groundwater affects more than 200 million people (3), and a global model places 350 million above the threshold for non-carcinogenic risk from drinking water alone (4). The resulting disease burden includes cancer of the lung, urinary bladder and skin, together with cardiovascular and neurodegenerative disease (3, 5, 6). The United States maximum contaminant level of 10 μg/l was set by treatment and analytical feasibility rather than by a demonstrated no-effect threshold (7). Diet is a second and less regulated route: rice concentrates iAs from soil and irrigation water (8), and surveys report up to 85 μg/kg iAs in infant rice cereals, delivered to a small body mass during a developmental window (9). Whether exposures in this range are genotoxic, and by what mechanism, remains contested (10, 11).

Exposure alone does not predict outcome. Among people with comparable intake, some develop skin lesions, cancer or cardiovascular disease and others do not, and the variance has been attributed to genetic background, DNA repair capacity, nutritional status and the gut microbiome (11, 12). Genetic mapping supports a host contribution: quantitative trait loci for arsenic-induced cellular phenotypes map across genetically diverse cells (13), and variant in the arsenic methyltransferase AS3MT associate with arsenic-induced skin lesions in exposed populations (14). Almost all of this work concerns metabolism, which acts on arsenic that has already crossed the plasma membrane. Uptake sits upstream. The dose that matters to a genome is the arsenite concentration reaching the cytoplasm, not the concentration in the water supply, and transporters set that quantity (15, 16). Population studies have begun to implicate the channels themselves, with AQP9 variant associating with urinary arsenic metabolite profiles in exposed women (17). Host genetic control of uptake would increase variation in internal dose across individuals exposed to the same external level, a gene-by-environment interaction proposed far more often than it has been tested. At physiological pH, trivalent arsenic exists largely as the uncharged species As(OH)_3_, close enough to glycerol in size and hydrogen-bonding geometry to enter cells through the aquaglyceroporin subfamily of aquaporins (18, 19). AQP7 and AQP9 conduct arsenite in Xenopus oocytes and in mammalian cells (18, 20), and the subfamily has been described more generally as a route for metalloid entry (16, 19). Aquaporin-3 (AQP3) belongs to the same subfamily and transports water, glycerol and hydrogen peroxide(21). Its status as an arsenite channel, however, is unresolved. Heterologous flux assays say no: the one side-by-side oocyte comparison of the four human aquaglyceroporins reported hAQP9 > hAQP7, with little or no As(OH)_3_ permeability for hAQP3 or hAQP10 (22), and a killifish AQP3 orthologue conducts water, glycerol and urea but not arsenite, the difference mapping to three carboxy-terminal residues (23). Human cell data indicate otherwise: an arsenite-resistant lung adenocarcinoma line has two-fold reduced AQP3, AQP3 knockdown lowers arsenic accumulation and confers resistance, and AQP3 overexpression in HEK293T cells raises both accumulation and As(III) susceptibility, with AQP7 and AQP9 barely detectable in either line (24); a genome-wide CRISPR screen in K562 cells identified loss of AQP3 as a determinant of arsenic trioxide tolerance (25); AQP3 and AQP10 participate in iAs uptake across human intestinal epithelium, where As(III) exposure upregulates both (26); and tissue arsenic retention tracks Aqp3 messenger RNA in mice (27).

AQP3 reaches this problem from a second direction. A functional screen for proteins whose overexpression raises endogenous DNA damage and mutation rate without disabling a repair pathway identified AQP3 as a candidate acting across human cell types (28, 29). Transcriptome-wide association analysis independently identified a lung adenocarcinoma susceptibility locus at 9p13.3 driven by higher predicted AQP3 expression (29), a locus carried forward in cross-ancestry meta-analysis of 61 047 lung cancer cases and 947 237 controls (30), in a disease whose inherited component extends from common susceptibility loci to rare deleterious germline variants (31); integration of lung tissue proteomics with genome-wide association data identifies AQP3 protein as one of two adenocarcinoma-specific susceptibility proteins (32). AQP3 is elevated in human arsenical skin cancers and in arsenic-treated keratinocytes (33). A transporter that is itself a lung cancer susceptibility gene, and that may admit a lung carcinogen, is worth testing directly.

Four gaps follow. First, the interaction has been asserted rather than tested: two independently acting genotoxins produce a larger combined effect without interacting, so a larger signal in channel-expressing, arsenite-treated cells is not evidence of synergy. The test is whether the gap between channel-expressing and control cells widens with dose. Second, whether any interaction is AQP3-specific or general to the subfamily is unknown: the paralogues have never been compared on a common host-cell background. Third, mutagenesis has not been measured at these doses. γH2AX reports breaks present at fixation, not heritable change; accurately repaired breaks leave no trace (34–36). Somatic mutation frequencies in normal human cells are 10⁻⁷ to 10⁻⁶ per base pair (37, 38), far below the error floor of short-read sequencing. Arsenic compounds this: as a weak direct mutagen and a potent co-mutagen (39–41), its mutational consequence need not track its damage signal. Duplex sequencing reaches that range (42, 43) and is validated for genotoxicity testing in human cells (44, 45). Fourth, the transcriptional response is heterogeneous, and bulk measurements average over it. Neither AQP3 expression nor arsenite response is uniform across cells, so a dose-dependent gap measured in bulk cannot distinguish a uniform shift in every cell from a change confined to a subpopulation. A gene-environment interaction of this kind is not a single response curve but a distribution of them.

Here we ask whether AQP3 expression changes the arsenite dose-response for DNA damage in human cells, whether it changes the mutational consequence of an exposure in the dietary range, and how the transcriptional response is distributed across cells. We show that the difference between AQP3-expressing and control cells widens with arsenite dose rather than adding a constant offset, that it requires the AQP3 sequence and is graded across point mutants, that N-acetylcysteine suppresses it in proportion to arsenite dose, that it depends on cell type, and that it is shared with AQP9 and AQP10. We then show that arsenite alone leaves somatic mutation frequency unchanged at doses up to 10 μM, while the mutational response to the same exposure differs between AQP3-positive and AQP3-negative cells sorted from one transfection. Finally, we use single-cell and ten-cell SHERRY transcriptomics to resolve how subpopulations shift under AQP3 expression and low-dose arsenite, separating changes confined to a subset of cells from shifts common to all.

## MATERIALS AND METHODS

### Cell lines and culture

MRC5-SV40 (SV40-immortalized human lung fibroblasts), HEK293T and RKO (BRAF wild type; a gift from the Yun laboratory) were maintained at 37°C in 5% CO_2_ and passaged before confluence in Dulbecco’s modified Eagle’s medium (DMEM; Invitrogen, 11965-092) supplemented with 10% fetal bovine serum, 2 mM L-glutamine, penicillin and streptomycin (Gibco). Lines were authenticated by short tandem repeat profiling (ATCC) and tested routinely for mycoplasma by sequencing.

### Plasmid constructs

Gateway entry clones for AQP3 were synthesized and cloned into pDONR plasmids. The point mutants H53F, W128A and S152A, previously characterized in human AQP3 for water permeability, pH sensitivity and Ni^2+^ inhibition (46), and the paralogues AQP7, AQP9 and AQP10, were synthesized in the same way. All clones were recombined into the N-terminal GFP destination vector pcDNA6.2/N-EmGFP-DEST using Gateway LR Clonase II (Thermo Fisher, 11791020), transformed into One Shot OmniMAX 2-T1R (Thermo Fisher, C854003), and verified by BsrGI-HF (New England Biolabs, R3575S) digestion and whole plasmid sequencing. LR reactions combined 50-150 ng of entry clone with 150 ng of destination vector in TE buffer, pH 8.0, to 8 μl, received 2 μl of LR Clonase II, were incubated at 25°C for 1 h and terminated with 1 μl of proteinase K (Thermo Fisher, EO0491) at 37°C for 10 min. Transformants were selected on LB agar with 100 μg/ml ampicillin (Sigma-Aldrich, A9518); plasmid

DNA was recovered with a QIAprep spin miniprep kit (Qiagen, 27106), digested with BsrGI-HF in CutSmart buffer (New England Biolabs, B6004S) for 1 h at 37°C, and resolved on 1% agarose in 1× TAE stained with SYBR Safe (Thermo Fisher, S33102) against a GeneRuler 1 kb ladder (Thermo Fisher, SM0311). GFP-Tubulin served as the transfection control throughout and mock-transfected cells as the gating control.

### Transfection, arsenite exposure and treatments

Cells were seeded at 5 × 10^5^ per well in six-well plates. Thirty minutes before transfection the medium was replaced with fresh serum-containing DMEM. For each well, 1 μg of plasmid DNA in 40 μl of serum-free DMEM and 2.5 μl of GenJet In Vitro DNA Transfection Reagent Ver. II (SignaGen, SL100488) in 40 μl of serum-free DMEM were held separately for 5 min, combined, held for a further 10 min at room temperature, and added dropwise. Medium was replaced the following day, and cells were harvested 72 h after transfection.

All arsenic exposures used trivalent inorganic arsenite, As (III), supplied as sodium arsenite (Ricca Chemical, # 714232) and added to the culture medium at 0, 1 or 5 μM for 72 h; duplex-sequencing exposures additionally included 10 μM. Arsenite was added after transfection and again at the same nominal concentration when the medium was replaced, so the concentration was restored once during the exposure. Doses are given throughout as the molar concentration of As (III) in the medium; where mass concentrations are quoted, they refer to elemental arsenic (74.92 g/mol) rather than to salt mass, so that 1 μM As (III) is ∼75 ppb and 5 μM is 375 ppb. Where indicated, N-acetylcysteine (6 mM) was added concurrently with arsenite at transfection and again at the medium change and maintained for the exposure. The AQP3 inhibitor DFP00173 (47) was used at 25 μM with DMSO as vehicle as described in the original study. N-ethyl-N-nitrosourea (ENU) was used at 400 and 2000 μM for 72 h as a positive control for mutagenesis.

### A mock-anchored damage threshold makes γH2AX percentages comparable across experiments

DNA damage was measured as phosphorylation of histone H2A.X at Ser139 (34) by indirect immunofluorescence on fixed, permeabilized intact cells read by flow cytometry(28–31), an endpoint validated across laboratories as a genotoxicity readout (36). Cells were harvested by trypsinization and ∼1 × 10^6^ cells per tube carried into staining. Cells were pelleted at 1000 × g for 5 min at 4°C, fixed in freshly prepared 2% (w/v) formaldehyde in PBS for 15 min on ice, permeabilized in 0.05% Triton X-100 for 15 min on ice, and blocked in 5% BSA/PBS (Cell Signaling Technology, 9998S) for 30 min on ice. Cells were stained with anti-phospho-histone H2A.X (Ser139) clone JBW301 (1:750; Sigma-Aldrich, 05-636-25UG) for 1 h at room temperature, washed, and incubated with goat anti-mouse IgG (H+L) Alexa Fluor 647 (1:1000; Invitrogen, A-21235) for 1 h at room temperature in the dark. Data were acquired on a Bio-Rad YETI/ZE5 flow cytometer. Every experiment included a mock-transfected, antibody-stained tube, which supplied that experiment’s gating threshold.

Because transfection is transient, every tube contains construct-expressing (GFP-positive) and untransfected (GFP-negative) cells that received the same dose, medium, staining and instrument settings (Supplementary Figure S2A-C). Gates were defined in FlowJo (Supplementary Figure S2A-C) and reproduced computationally for analysis following the approach used previously in this laboratory (28); reproduced populations matched FlowJo’s own event counts with minimal discrepancy (Supplementary Figure S2A-C). Events were gated on forward and side scatter, then on FSC-A/FSC-H for single cells, then split into GFP-positive and GFP-negative populations using a fixed intensity threshold set from the mock-transfected control of that experiment.

For each experiment the γH2AX threshold was set at the top 0.5% of the mock-transfected, antibody-stained single-cell distribution and applied unchanged to every tube acquired that day (Supplementary Figure S2D). The threshold is a within-experiment normalization rather than a biological baseline: because it is re-derived on each acquisition date, percentages above threshold are comparable across days whereas absolute fluorescence is not. All conclusions were re-tested with the threshold at the top 1%, 2% and 5% (Supplementary Figure S2D-H) (Figure 1C, F). The primary readout is the percentage of GFP-positive cells above threshold. Two secondary readouts are reported: the within-tube contrast, defined as the percentage above threshold in GFP-positive minus GFP-negative cells of the same tube; and the median γH2AX intensity within the GFP-positive gate, normalized to GFP-Tubulin.

**Figure 1.**
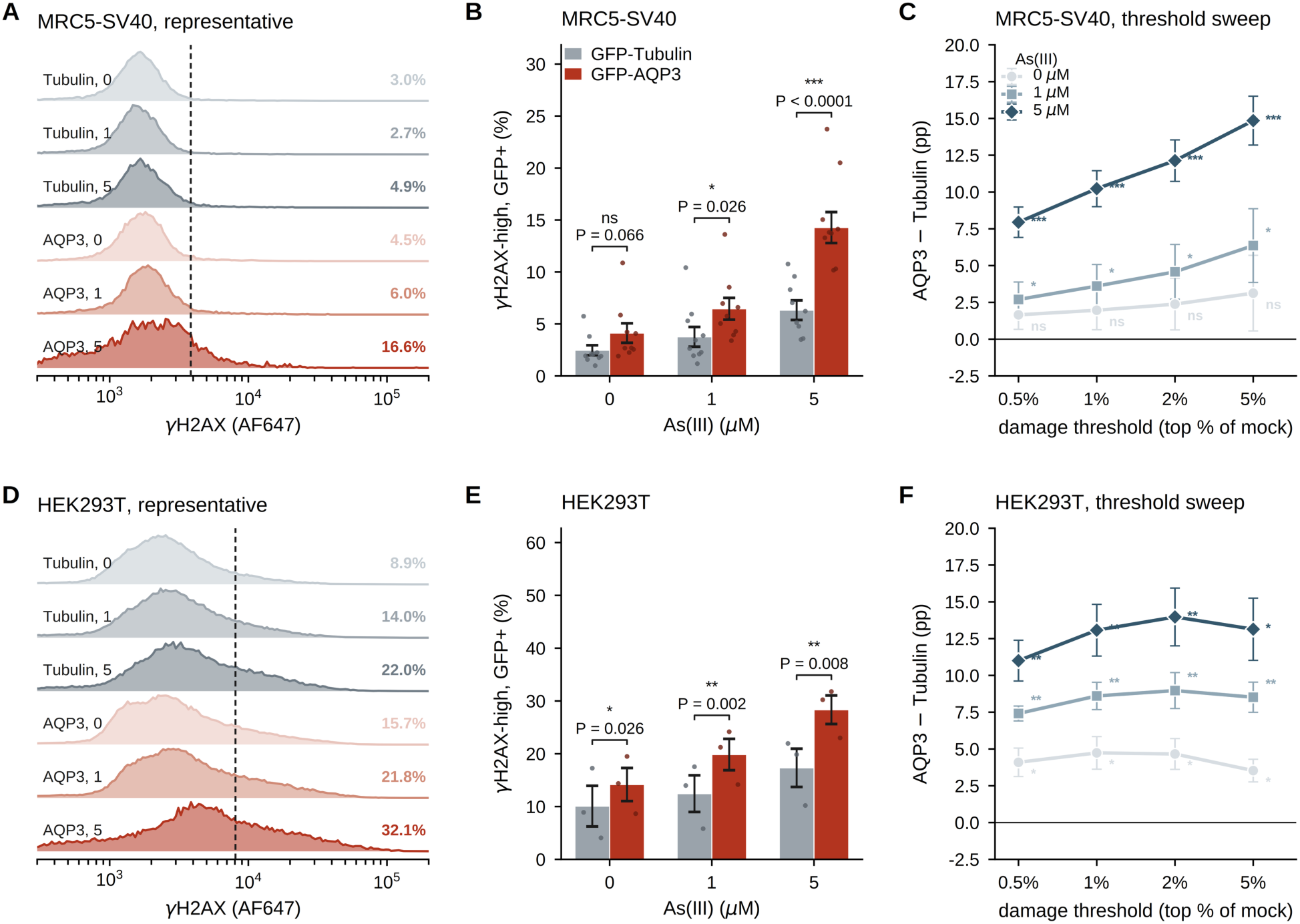
AQP3 makes arsenite genotoxicity dose-dependent. One row per cell line: MRC5-SV40 (A-C) and HEK293T (D-F). (A, D) Representative γH2AX distributions in GFP-positive cells of one acquisition, GFP-Tubulin and GFP-AQP3 at each dose; dashed line, that plate’s damage threshold; percentage, γH2AX-high in the tube. (B, E) γH2AX-high across 0, 1 and 5 μM As(III); bars, mean ± SEM across acquisition dates (MRC5-SV40 n = 9, 9, 8; HEK293T n = 3); points, individual dates; brackets, one-sided paired t-test against the date-matched GFP-Tubulin control. (C, F) The AQP3-minus-Tubulin difference at each dose recomputed across all four damage thresholds, from the top 0.5% to the top 5% of the mock distribution; mean ± SEM over the same dates. In MRC5-SV40 the contrast is significant at every threshold at 1 and 5 μM and at none without arsenite (P = 0.066 to 0.129), which is what the synergy of Figure 2 predicts. In HEK293T it is significant at every threshold at every dose, including without arsenite (P = 0.022 to 0.026), so in that line AQP3 raises γH2AX by roughly 4 pp on its own before any arsenite is added.

### Technical controls on the flow cytometry dataset

Transfection efficiency did not fall with arsenite: across transfected tubes in MRC5-SV40 the GFP-positive fraction was 18.3%, 18.7% and 17.8% at 0, 1 and 5 μM (55-65% in HEK293T, 7-10% in RKO), so the populations being compared are of similar size across doses (Supplementary Figure S3). Cell recovery did fall. The fraction of acquired events passing the singlet gate declined from 73.8% to 71.3% to 64.2% across 0, 1 and 5 μM in MRC5-SV40, with comparable declines in HEK293T (64.1%, 59.9%, 55.1%) and a smaller one in RKO (86.4% to 81.9%). At 1 μM this loss is negligible; at 5 μM it is not, and because apoptotic cells generate pan-nuclear γH2AX, every 5 μM contrast is read as an upper bound on repairable damage. Critically, the loss is the same in the two arms being compared: at 5 μM in MRC5-SV40, singlet recovery was 63.5% in AQP3-transfected and 63.6% in GFP-Tubulin-transfected tubes, and the GFP-positive fraction 18.4% and 19.9%, so differential cell loss does not generate the AQP3-versus-Tubulin difference.

### Fluorescence-activated cell sorting

MRC5-SV40 cells were transfected with pcDNA6.2/N-EmGFP-AQP3 or mock-transfected, with As (III) added at the time of transfection and maintained for 72 h, matching the γH2AX exposure. At the end of the exposure, cells were sorted on a Bio-Rad S3e Cell Sorter (Bio-Rad Laboratories, Hercules, CA, USA) under ProSort software control, with EmGFP excited by the 488 nm laser and detected in FL1 (525/30 nm bandpass), into GFP-positive (AQP3-overexpressing) and GFP-negative (AQP3-negative) fractions taken from the same transfected population. The two genotypes compared within a sequencing run therefore share transfection, exposure, sorting, passage, culture history, library batch and sequencing run. Sorted samples cover As (III) at 0 and 1 μM only. Runs designated parental used unsorted MRC5-SV40 cultures exposed to As (III) at 0, 1, 5 or 10 μM, or to ENU at 400 or 2000 μM; the 5 and 10 μM As (III) exposures and all ENU exposures were unsorted. The untransfected status of parental cultures is recorded by the absence of a transgene annotation rather than positively.

### Duplex sequencing library preparation, sequencing and analysis

Genomic DNA was extracted with a column-based blood and tissue kit (Qiagen), quantified fluorometrically and sonicated on a Bioruptor system (Diagenode) to a median fragment size of 300-400 bp, with TapeStation 4200 average fragment sizes of 302-422 bp recorded for the sonicated aliquots.

Libraries were prepared with TwinStrand DuplexSeq adapters carrying an inline dual unique molecular identifier on both reads. Two capture panels were used: the TwinStrand DuplexSeq human mutagenesis panel human-muta-v1.0, comprising 20 targets (48 kb total), validated for human-cell mutagenesis measurement(48); and a custom ∼200 kb panel tiling the EGFR locus, target interval chr7:55,018,933-55,211,713 (GRCh38/hs38DH), with 120 bp biotinylated capture baits (Integrated DNA Technologies) tiling introns and exons and recovered on streptavidin magnetic beads. The intended input was 250 ng of sheared genomic DNA per library; two libraries did not reach it because template concentration was limiting and the entire eluate was carried forward instead, and low genome equivalents inflate duplex mutation-frequency estimates. Twenty-six libraries were sequenced paired-end on NextSeq 550 or NextSeq 2000 instruments and demultiplexed with bcl2fastq or bcl-convert.

Reads were processed with a locally implemented duplex-consensus pipeline rather than the vendor’s hosted application, which was discontinued during this work. FASTQ files were converted with fgbio FastqToBam, aligned with bwa mem, grouped by unique molecular identifier under the paired dual-UMI strategy, called to single-strand and then duplex consensus with CallDuplexConsensusReads and FilterConsensusReads, realigned, hard-clipped across read overlaps, and called with VarDictJava (49) configured to retain single-duplex-molecule calls. Alignment was to GRCh38 in the hs38DH configuration; the decoy-containing build is required because decoy-derived mismapping into a small capture panel is not negligible when calling at 10^-7^. The implementation was validated against the vendor application on two libraries carried through both: germline coordinate concordance was 48/48 and 54/54, six-channel spectral cosine similarity 0.9952 and 0.9736, and the mutation frequency fell inside the vendor application’s Poisson 95% confidence interval in both cases (P = 0.354 and 0.182).

Calls were classified by variant allele fraction (VAF) as germline at ≥0.30, clonal or subclonal at 0.01-0.30 (excluded), and somatic at <0.01; mutations supported by a single duplex consensus molecule were retained by design, and only single-nucleotide variants were counted. Three filters were then applied uniformly: a position filter requiring the mean position of the variant within its supporting reads to be at least 8 bases from the rear end (VarDict PMEAN ≥ 8), swept at 0, 5, 8 and 10 without reprocessing; a recurrent-position blacklist excluding any somatic key called in three or more independent libraries on the same panel, rebuilt at each PMEAN setting; and a per-read-length mappability mask restricting all samples within a comparison to territory uniquely mappable at every read length involved (50). A region-level mask excluding a 5 kb misalignment window on the EGFR panel (chr7:55 163 933-55 168 933) was applied as a sensitivity analysis and to the binned EGFR display, and every contrast is reported with and without it. Mutation frequency was calculated as the number of distinct somatic mutations divided by the number of duplex base pairs interrogated (MF_min_).

#### Low-input RNA sequencing (SHERRY)

Libraries were prepared by SHERRY, in which Tn5 tagments an RNA/DNA hybrid directly (51). The protocol was first optimized on HEK293T cells across RNA inputs of 10, 30 and 300 pg and reverse-transcriptase amounts of 10, 25, 50 and 100 U per reaction, with additional Tn5-amount, Bst-concentration and single-cell variants; these 30 technical libraries established the working conditions and were not used for any biological analysis. The reverse-transcriptase titration was a complete 3 × 4 factorial (300, 30 and 10 pg RNA × 10, 25, 50 and 100 U, one library per condition with Tn5 held constant), scored on median insert size, exonic assignment rate, alignments per fragment, duplicate fraction and genes detected after rarefaction to a common depth of 200,000 assigned reads (Supplementary Figure S4). Twenty-five units was optimal at 300 pg on every metric, whereas at 10 and 30 pg depth-matched complexity was still rising at the top of the range, so 100 U was used for the single-cell and ten-cell libraries; with one library per condition these are directional observations rather than tested differences.

Single cells and pools of ten cells were sorted as described above into low-profile tubes (Bio-Rad, TLS0801 and TCS0803) containing 3.45 µl of lysis buffer (1 U RNase inhibitor, Invitrogen AM2696; 1 µl of 10 µM oligo-dT primer; 1 µl of 10 mM dNTPs; 0.5% Triton X-100), and lysed at 65 °C for 10 min. Sorted libraries provide the AQP3-positive versus GFP-negative comparison at 0 and 1 µM As (III); unsorted parental cultures provide the dose ladder.

Reverse transcription used SuperScript IV (Invitrogen, 18090010) in 6.55 µl with 5× first-strand buffer, 8 mM DTT, 2.5 M betaine (Sigma-Aldrich) and RNase inhibitor, at 50 °C for 50 min then 80 °C for 10 min. The hybrid was tagmented at 55 °C for 30 min in 20.5 µl of 4× TD buffer with Tn5 transposase (Diagenode, C01070012-200), 16% PEG 8000, 0.5 mM ATP (New England Biolabs, P0756L) and RNase inhibitor. Gaps were filled with Q5 High-Fidelity master mix (New England Biolabs, M0492S) and Bst 3.0 (New England Biolabs, M0374L) at 72 °C for 15 min and 80 °C for 10 min. Libraries were indexed by PCR with Q5 High-Fidelity 2× master mix and 10 µM Nextera i5 and i7 index primers (98 °C for 30 s; 24 cycles of 98 °C for 20 s, 60 °C for 20 s, 72 °C for 2 min; 72 °C for 5 min), checked on a TapeStation 4200 and sequenced on a NextSeq 2000 with P2 or P3 100-cycle kits.

#### SHERRY analysis

Reads were aligned with nf-core/rnaseq (52) (STAR 2.7.11b, GRCh38 Ensembl 109) and counted with featureCounts in unstranded mode; the exonic assignment rate of each library was taken from the featureCounts summary as reads assigned to exons divided by all counted reads. Differential expression and the AQP3-by-arsenite interaction were fitted in DESeq2 with median-of-ratios size factors and sequencing run as a covariate. Genes were tested only if they carried at least five counts in at least eight of the analyzed ten-cell libraries, leaving 120 of the 124 panel genes measurable. Because the interaction statistic correlates with expression level, gene-set tests compared the observed set median of the interaction statistic with 5,000–10,000 random gene sets matched on baseMean ventile. The combined condition was additionally contrasted against each single condition within the one sequencing run that contained all four groups of the design; a gene was called above both singles only if it reached a 5% false-discovery rate in both contrasts with a positive fold change in each.

### Statistics

Replicate tubes were averaged within an acquisition date to give one value per construct, dose and date, and each construct was compared with the GFP-Tubulin control acquired on that same date. Every flow cytometry test is therefore paired within date, which removes between-day variation in staining, instrument setting and transfection efficiency, and n denotes the number of independent acquisition dates rather than the number of tubes. The hypothesis is directional and was specified in advance (an aquaglyceroporin raises γH2AX and N-acetylcysteine lowers it), so every such contrast is tested by one-sided paired t-test on the paired difference at α = 0.05, applied uniformly to all of them. Intervals are two-sided 90%, whose lower bound is the one-sided 95% bound, so interval and P value always agree; the two-sided, log-ratio and Wilcoxon value of every contrast is tabulated in the source data. Controls and exploratory analyses (cell recovery, the DFP00173 arm and transfection efficiency) have no pre-specified direction and remain two-sided. Point estimates are the mean paired difference. The pairing unit is the acquisition date for the AQP3-versus-GFP-Tubulin arsenite dose series, and the acquisition plate for the paralogue, RKO and N-acetylcysteine comparisons, because several dates carry more than one plate and date-level pairing would admit control tubes from a plate that did not carry the construct. Where a plate carries two tubes per arm those tubes are averaged within the plate before testing; a linear mixed model with plate as a random intercept, tested on degrees of freedom, is reported as a secondary analysis and leaves every point estimate unchanged. Because the control baseline differs several-fold between cell lines, percentage-point differences are not directly comparable across lines and the construct-to-control ratio is reported alongside the difference wherever lines are compared. Contrasts resting on two experiment dates are reported descriptively, with both individual values given, and carry no P value: a paired t-test on two experiments has one degree of freedom and its P value is determined by how closely two replicates happen to agree rather than by the size of the effect.

The interaction was tested as the 2 × 2 factorial within an acquisition date (neither factor, gene alone, environment alone, both) by taking the excess of the observed combination over the sum of the two single increments, which tests an additive (Bliss) null on the percentage-point scale and equals the construct × dose interaction term. Mixed-model estimators use all tubes with acquisition date as a random intercept and are tested on degrees of freedom, because the contrast is a between-date quantity however many tubes were run. One primary contrast was specified in advance: MRC5-SV40, top 0.5% threshold, AQP3 wild type against GFP-Tubulin across 0, 1 and 5 μM together with the dose-step interaction. All other contrasts are exploratory.

For the duplex-sequencing contrasts a rate ratio was formed from two mutation frequencies and the variance of its logarithm taken as the sum of the reciprocals of the two mutation counts (Poisson); genotype comparisons are ratios of ratios formed within a single sequencing run, and contrasts were pooled by fixed-effect inverse-variance weighting on the log scale. Contrasts that share a denominator are correlated, and the genotype comparisons do: two of the three are different AQP3-positive pairs measured against one AQP3-negative pair. Those were pooled by generalized least squares with the covariance between two such contrasts set to the variance of the shared log ratio; I2 is not defined for that pool and is not reported with it. That variance model treats every interrogated duplex base as an independent trial, so it understates uncertainty relative to a model in which the library is the replicate; where the two disagree materially, both are reported.

### Software, randomization

Gate reproduction, quantification and all statistical analysis were performed in Python 3 with NumPy, SciPy and pandas. Acquisition order was not randomized, and the analyst was not blinded to condition. Gating and thresholding were governed by one rule applied identically to every tube in an experiment, and all quantification was executed by script from the raw event data rather than by manual reading.

## RESULTS

### AQP3 sensitizes inorganic arsenic induced genotoxicity

The question this study asks is whether the channel complement of a cell changes what a given external arsenite exposure does to its genome. We therefore held the exposure fixed and varied the transporter, expressing GFP-tagged AQP3, three AQP3 point mutants and the paralogues AQP7, AQP9 and AQP10 against a GFP-Tubulin control in three human cell lines at 0, 1 and 5 μM As (III) for 72 h. Three readouts were applied to the same system, each resolving a different consequence and a different unit: γH2AX by flow cytometry counts how many cells carry breaks at the moment of fixation; duplex sequencing measures heritable single-nucleotide change at 10^-7^ per base pair; and SHERRY, a low-input RNA method applied to pools of ten sorted cells, reports transcriptional state at a grain fine enough to show whether a response is shared by all cells or confined to a few.

### A mock-anchored threshold makes damage comparable across experiments

Comparing a transfected construct with a control across many acquisition days requires a damage scale that does not drift with staining or instrument setting. We therefore scored damage as the percentage of construct-expressing cells whose γH2AX signal exceeded a threshold taken from the mock-transfected, antibody-stained cells of the same experiment, at the top 0.5% of that distribution (Supplementary Figure S2A-D). Because the threshold is re-derived each day, percentages are comparable across days while absolute fluorescence is not; because transfection is transient, each tube also carries its own untransfected population as an internal control. On a representative acquisition the readout separates the conditions cleanly: 2.9% of GFP-Tubulin cells exceeded the threshold without arsenite and 4.8% at 5 μM, against 4.3% and 15.8% for GFP-AQP3 (Supplementary Figure S2E-H).

Before any aquaglyceroporin was introduced, arsenite raised the γH2AX-high fraction with dose in two of the three lines. In GFP-Tubulin-transfected MRC5-SV40 lung fibroblasts the fraction ran 2.47%, 3.76% and 6.33% across 0, 1 and 5 μM on the dates carrying the AQP3 comparison, and in HEK293T cells 10.08%, 12.44% and 17.34%. RKO colorectal carcinoma cells showed no dose-response over the same range (6.49%, 6.86%, 6.38%). The two responsive lines therefore bracket an exposure range in which a transporter effect can be detected, while RKO supplies a background in which arsenite alone does nothing measurable.

HEK293T serves throughout as the technical evaluator for the assay. Transfection efficiency there has a median of 65% of single cells against 22% in MRC5-SV40 and 12% in RKO (Supplementary Figure S3A), so the construct-expressing population being scored is three-to sevenfold larger and per-tube sampling error correspondingly smaller. Every qualitative feature of the MRC5-SV40 result reproduces in that setting: AQP3 shifts the distribution rightward at 5 μM where GFP-Tubulin shifts less (Figure 1D); the same holds for the paralogues and for the H53F mutant in that line, shown with their own sections below.

### The AQP3 effect is a gene-by-dose interaction, not an additive offset

AQP3 raised the γH2AX-high fraction at every dose in MRC5-SV40 cells (Figure 1A, B). Without arsenite the fraction was 4.13% against 2.47% for the date-matched control (+1.65 percentage points, 90% CI −0.18 to +3.49; P = 0.066, n = 9). Fold changes are the geometric mean of the within-date ratios, which is 1.62-fold here and need not equal the ratio of the two means. At 1 μM it was 6.46% against 3.76% (+2.70 points, 90% CI +0.50 to +4.90, 1.93-fold; P = 0.026, n = 9), and at 5 μM 14.27% against 6.33% (+7.95 points, 90% CI +5.99 to +9.91, 2.36-fold; P = 0.0001, n = 8). The contrast is significant at 1 and 5 μM and not without arsenite, and that pattern holds at all four damage thresholds (P = 0.066 to 0.129 without arsenite; Figure 1C), which is what the synergy of Figure 2 predicts. One micromolar arsenite corresponds to 74.9 ppb elemental arsenic, an environmentally relevant concentration, and 5 μM to 375 ppb, both as free concentrations sustained in culture medium over 72 h.

**Figure 2.**
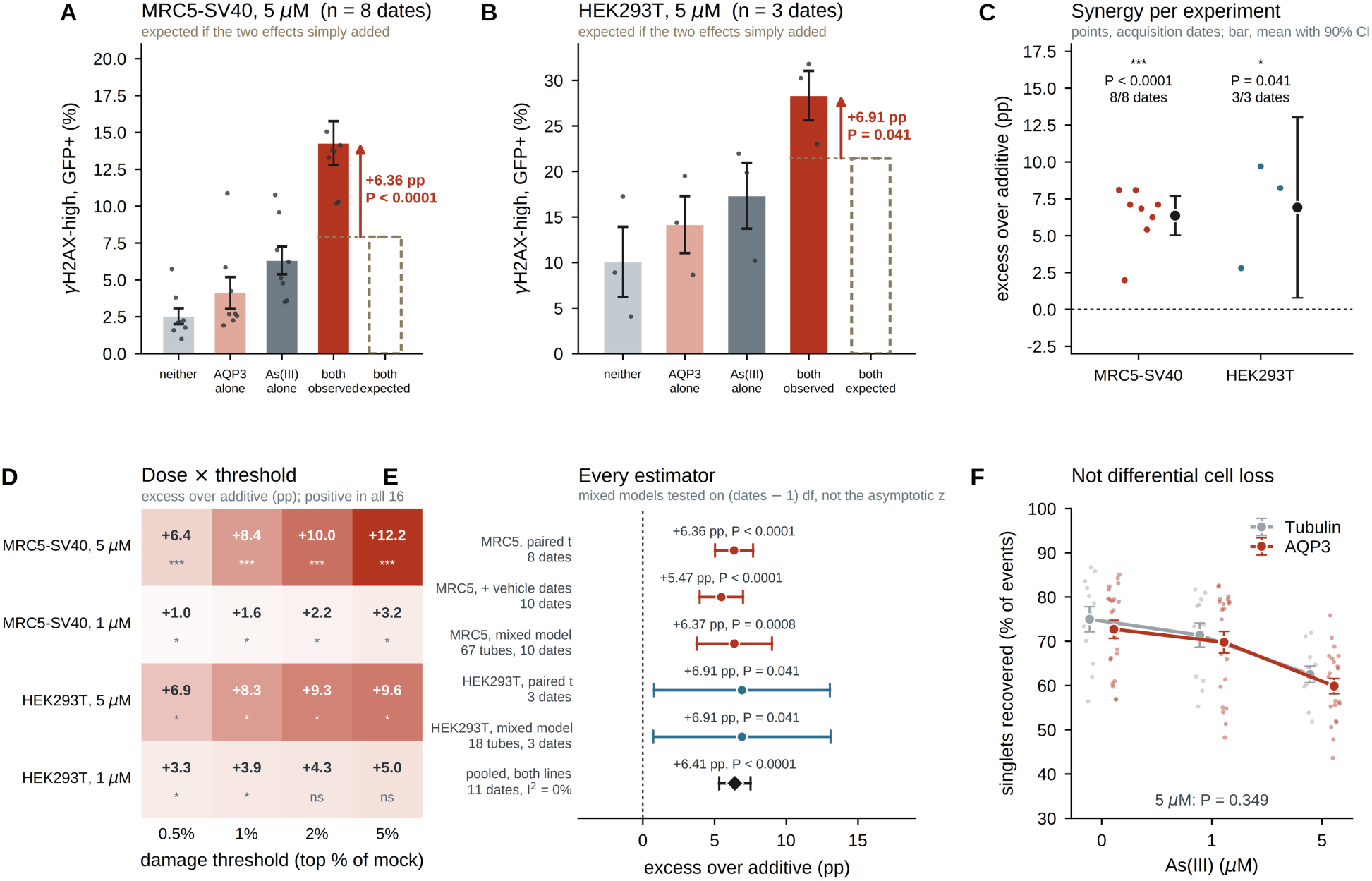
AQP3 and arsenite act synergistically, not additively. The 2 × 2 factorial, taken within each acquisition date: neither factor (GFP-Tubulin, 0 μM), gene alone (GFP-AQP3, 0 μM), environment alone (GFP-Tubulin, 5 μM), and both (GFP-AQP3, 5 μM). If the two effects simply added, the combination would sit at (gene alone − neither) + (environment alone − neither) above baseline; the excess over that expectation is the synergy, and equals the construct × dose interaction term. (A) MRC5-SV40: AQP3 alone +1.59 pp, 5 μM As(III) alone +3.78 pp, additive expectation +5.37 pp, combination +11.73 pp, an excess of +6.36 pp (90% CI +5.03 to +7.68, P < 0.0001), more than double what additivity predicts. (B) HEK293T: expectation +11.35 pp, observed +18.26 pp, excess +6.91 pp (90% CI +0.78 to +13.04, P = 0.041) on three dates. Bars, mean ± SEM across dates; points, individual dates; dashed outline, the additive expectation. (C) The excess per acquisition date: positive on 8 of 8 MRC5-SV40 and 3 of 3 HEK293T dates. (D) The excess at both arsenite doses and all four damage thresholds: positive in all sixteen combinations and significant in fourteen. (E) The same quantity under every estimator. The mixed models use all tubes with acquisition date as a random intercept and are tested on (dates − 1) degrees of freedom rather than the asymptotic z, because the contrast is a between-date quantity however many tubes were run. All five estimates fall between +5.47 and +6.91 pp, and the two lines pool to +6.41 pp (90% CI +5.32 to +7.51) with I² = 0%. (F) Singlets recovered as a fraction of recorded events; the fall at 5 μM is the same in both arms (P = 0.35, two-sided), so the excess is not produced by differential cell loss. Synergy is defined here on the percentage-point (Bliss) scale. On the multiplicative scale the combination exceeds the product of the single effects by 1.51× in MRC5-SV40 (P = 0.056). The combination also exceeds each single agent alone in both lines (MRC5-SV40 P < 0.0001 against either).

The pattern of three dose-wise contrasts is not itself evidence of an interaction, because two agents that each damage DNA independently produce a larger combined effect without interacting. We therefore read the experiment as the 2 × 2 factorial it is, taken within each acquisition date: neither factor, gene alone, environment alone, and both. If the two effects simply added, the combination would sit at the sum of the two single increments above baseline, and the excess over that expectation is the interaction term (Figure 2A). In MRC5-SV40, AQP3 alone raised the γH2AX-high fraction by 1.59 percentage points and 5 μM arsenite alone by 3.78, so additivity predicts +5.37; the combination reached +11.73, an excess of +6.36 points (90% CI +5.03 to +7.68; P < 0.0001), more than double what additivity predicts. The excess was positive on all eight MRC5-SV40 dates and on all three HEK293T dates (Figure 2C), and positive at both arsenite doses across all four damage thresholds: sixteen combinations, significant in fourteen (Figure 2D). Five estimators, including mixed models fitted to every tube with acquisition date as a random intercept and tested on degrees of freedom, fall between +5.47 and +6.91 points, and the two cell lines pool to +6.41 points (90% CI +5.32 to +7.51) with I2 = 0% (Figure 2E). AQP3 therefore changes the slope of the arsenite dose-response rather than adding a constant number of percentage points. Differential cell loss does not produce it: singlet recovery falls with dose by the same amount in both arms (AQP3 72.7% to 59.9%, GFP-Tubulin 75.0% to 62.5%; P = 0.35, two-sided; Figure 2F).

Synergy is defined here on the percentage-point (Bliss) scale, on which the null was pre-specified. The effect also grows on the ratio scale: AQP3 raised the γH2AX-high fraction 1.62-fold without arsenite, 1.93-fold at 1 μM and 2.36-fold at 5 μM. Against a multiplicative null the combination exceeds the product of the two single effects by 1.51-fold in MRC5-SV40, short of significance at this n (P = 0.056), and it exceeds each single agent alone in both lines (P < 0.0001). The interaction is therefore established on the scale on which it was specified and is in the same direction, without reaching significance, on the other.

The interaction was retained in a second host-cell background. In HEK293T cells AQP3 raised the γH2AX-high fraction by 4.09 percentage points without arsenite (90% CI +1.27 to +6.92; P = 0.026), by 7.41 points at 1 μM (P = 0.0023) and by 11.00 points at 5 μM (P = 0.0078; n = 3; Figure 1E), significantly at every dose and at every threshold (Figure 1F); the same 2 × 2 analysis gives an excess over additivity of +6.91 points (90% CI +0.78 to +13.04; P = 0.041), almost exactly the MRC5-SV40 value (Figure 2B). An AQP3 effect of that size in the complete absence of arsenite is not what MRC5-SV40 shows, and HEK293T is best described as carrying an arsenite-independent AQP3 component with a dose-dependent increment on top of it. A published precedent exists for an AQP3 transport phenotype in this line, where AQP3 overexpression raised arsenic accumulation and As (III) susceptibility and AQP3 knockdown lowered both (24).

Three controls place the primary measurement. Scoring each tube against its own GFP-negative population, which removes between-day variation entirely, gave an AQP3 excess of 3.54, 5.51 and 10.82 percentage points at 0, 1 and 5 μM against 1.93, 2.74 and 4.09 points for GFP-Tubulin on the same dates (n = 9, 9 and 8), so untransfected neighbours in the same well do not carry the effect (source data). Moving the damage threshold from the top 0.5% to the top 1%, 2% and 5% of the mock distribution widened the AQP3-minus-Tubulin difference monotonically, from +7.95 to +14.85 percentage points at 5 μM and from +2.70 to +6.36 at 1 μM, so no conclusion depends on one arbitrary cutoff (Figure 1C, F); these four gates are nested subsets of one measurement and are not four independent confirmations. Transfection efficiency does not carry the effect either: within an acquisition plate GFP-AQP3 exceeded GFP-Tubulin by +1.1 points in MRC5-SV40 (n = 11, P = 0.34), +0.6 points in HEK293T (n = 3, P = 0.84) and +3.1 points in RKO (n = 5, P = 0.50), none distinguishable from zero, and the size of the 5 μM excess tracked neither how well a plate transfected (r = +0.34, −0.38, −0.39) nor how unbalanced its two arms were (r = +0.43, +0.79, +0.69), with no consistent sign across lines (Supplementary Figure S3).

### The response requires His53, partly requires Trp128, and does not require Ser152

To determine whether the response depends on the AQP3 protein sequence rather than on the transfection of a membrane protein, we compared wild-type AQP3 with three point mutants previously characterized in human AQP3 (46) across the arsenite dose series (Figure 3A). On an MRC5-SV40 acquisition carrying all five constructs, wild-type AQP3, S152A and W128A track each other across the series (2.5 → 5.3 → 13.5%, 3.3 → 5.3 → 13.5% and 2.4 → 5.8 → 12.4%) while H53F stays with GFP-Tubulin (1.0 → 2.7 → 5.0% against 2.0 → 1.8 → 4.9%; Figure 3B-D). Placing each mutant between wild-type activity and the GFP-Tubulin level that marks complete loss, S152A sits on the wild-type line (83% of wild-type activity retained at 5 μM; −1.31 points, P = 0.30), W128A sits between the two references (53% retained; −3.69 points, P = 0.018) and H53F sits on the GFP-Tubulin bar (7% retained; −7.28 points, P = 0.0053) (Figure 3E). The His53 deficit holds in both cell lines and at every dose: −2.27, −3.45 and −7.28 points in MRC5-SV40 (P = 0.012, 0.0032, 0.0053) and −9.19, −9.50 and −12.08 points in HEK293T (P = 0.033, 0.028, 0.022) (Figure 3F). These estimates come from a linear mixed model fitted to every tube of both constructs at that dose with acquisition date as a random intercept and tested on (dates − 1) degrees of freedom, which uses all replicates without treating tubes from one plate as independent experiments; 54-55% of the variance in these contrasts is plate-to-plate, and each rests on 26-33 tubes over 10 acquisition dates in MRC5-SV40 and 9 tubes over 3 dates in HEK293T. The requirement is therefore graded rather than all-or-none and centers on His53, in the same construct backbone transfected the same way, so the effect follows the AQP3 sequence and not the act of loading the membrane with a GFP-tagged membrane-spanning protein.

**Figure 3.**
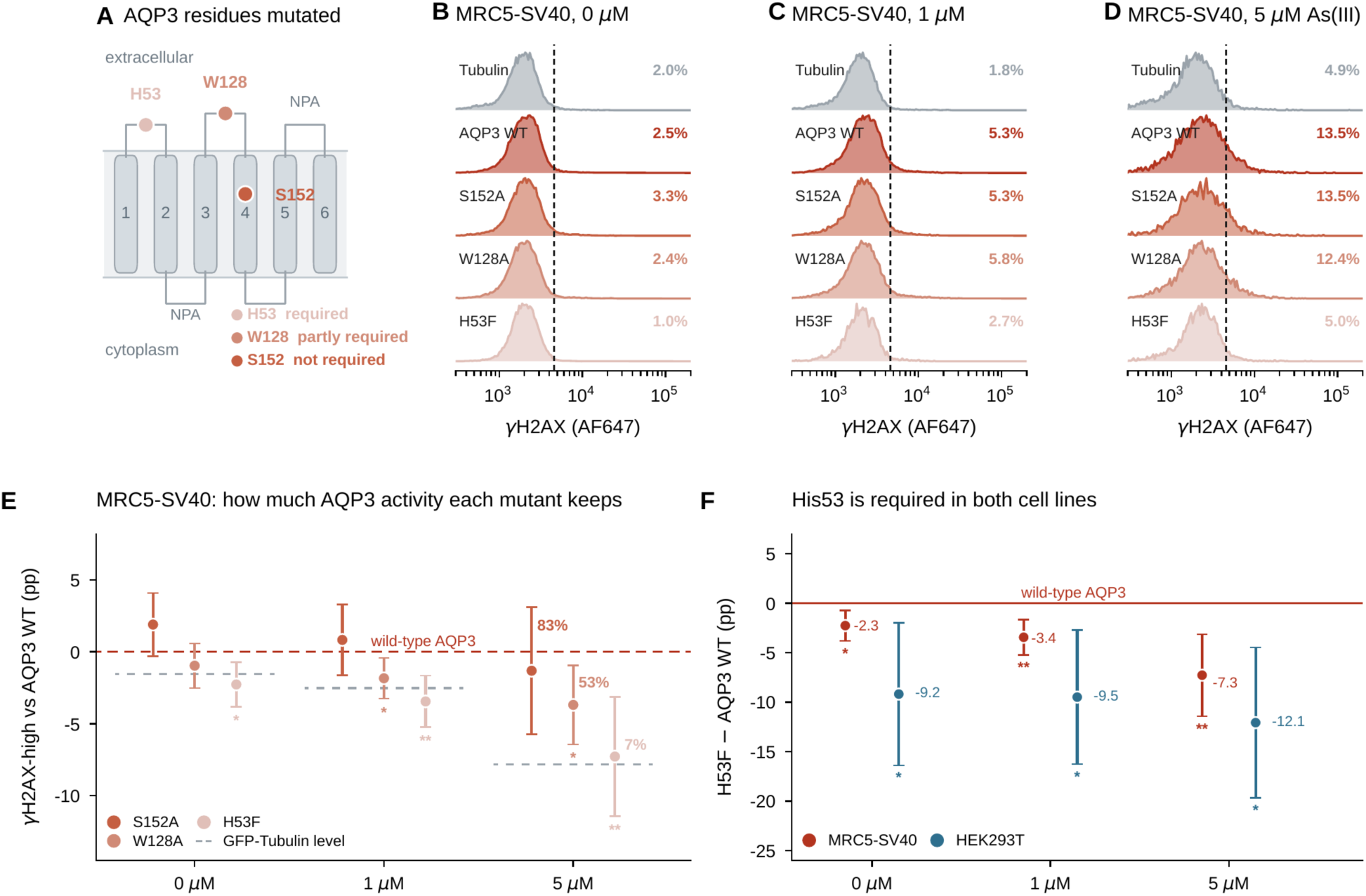
The AQP3 response requires His53, partly requires Trp128, and does not require Ser152. (A) The three mutated residues on the aquaporin six-transmembrane topology, with the verdict from E and F. (B–D) Representative γH2AX distributions in GFP-positive cells of one MRC5-SV40 acquisition carrying all five constructs (1 September 2023) at 0, 1 and 5 μM As(III); dashed line, that plate’s damage threshold; percentage, γH2AX-high in the tube. Wild-type AQP3, S152A and W128A track each other across the dose series (2.5 → 5.3 → 13.5%, 3.3 → 5.3 → 13.5% and 2.4 → 5.8 → 12.4%) while H53F stays with GFP-Tubulin (1.0 → 2.7 → 5.0% against 2.0 → 1.8 → 4.9%). (E) Each mutant placed between wild-type activity (red line, zero) and the GFP-Tubulin level that marks complete loss (grey bar); percentages at 5 μM are the fraction of wild-type activity retained. S152A sits on the wild-type line (83% retained, −1.31 pp, P = 0.30), W128A sits between the two references (53% retained, −3.69 pp, P = 0.018) and H53F sits on the GFP-Tubulin bar (7% retained, −7.28 pp, P = 0.0053). (F) The His53 deficit in both cell lines: below wild type in all six line × dose comparisons and significantly so in every one: MRC5-SV40 −2.27, −3.45 and −7.28 pp (P = 0.012, 0.0032, 0.0053) and HEK293T −9.19, −9.50 and −12.08 pp (P = 0.033, 0.028, 0.022). Estimates in E and F come from a linear mixed model fitted to every tube of both constructs at that dose, with acquisition date as a random intercept: this uses all replicates, including several from the same day, without treating tubes from one plate as independent experiments. The model separates plate-to-plate from tube-to-tube variance, and 54–55% of the variance in these contrasts is plate-to-plate, so the contrast is tested on (dates − 1) degrees of freedom. Intervals are 90%, matching the one-sided tests used throughout; mutant-versus-wild-type contrasts are one-sided because loss of function is directional. 26–33 tubes over 10 acquisition dates per contrast in MRC5-SV40, 9 tubes over 3 dates in HEK293T. The residues were characterized for water permeability, pH sensitivity and Ni²⁺ inhibition rather than for arsenite, and membrane localization of the mutants was not confirmed here, so a reduced response is a loss of detectable activity and not a demonstration that the channel is transport-dead.

These AQP3 mutants were characterized for water permeability, pH sensitivity and Ni^2+^ inhibition (46), not directly for arsenite; arsenite and water permeability are separable properties of AQP3, since in killifish AQP3 arsenite permeability is conferred by carboxy-terminal residues and is dissociable from water and glycerol transport (23). Because the mutants have not been shown to reach the plasma membrane, transport-deficient and mislocalized remain indistinguishable here. The sequence dependence is established; its structural basis is not.

### DNA damage tracks how much channel a cell carries

Within one tube every transfected cell carries a different amount of construct, read out as GFP intensity. Splitting the GFP-positive cells of a tube into five equal-count intensity bins turns that into a dose-response in expression level measured inside a single well: brighter and dimmer cells sat in the same tube, saw the same arsenite and the same stain, so there is no plate-to-plate normalization and no transfection-efficiency confound (Figure 4A-C). Damage rose with construct level for AQP3 at every arsenite concentration, and the AQP3 gradient was steeper than the GFP-Tubulin gradient throughout: the AQP3-minus-Tubulin excess across the bins grew from +17.9 to +29.4 percentage points across 0, 1 and 5 μM in HEK293T (P = 0.013, 0.010, 0.0016; Figure 4D-F) and from +4.8 to +8.5 points in MRC5-SV40 (P = 0.034, 0.0037, 0.0008; Figure 4G-I). Two features of the control matter for interpretation. GFP-Tubulin carries a gradient of its own, +8.0 points in MRC5-SV40 and +17.9 in HEK293T, so brightness is not inert, because more plasmid means more stress whatever it encodes, which is why the AQP3-minus-Tubulin excess rather than the raw AQP3 gradient is the channel-attributable quantity. And that excess is already present without arsenite, +17.9 points in HEK293T and +4.8 in MRC5-SV40, so it is not purely arsenite-dependent; what arsenite does is make it larger, by 1.6-fold in HEK293T and 1.8-fold in MRC5-SV40 from 0 to 5 μM. This is a dose-response in channel level that no difference in transfection efficiency can produce, because it is read between cells of one well.

**Figure 4.**
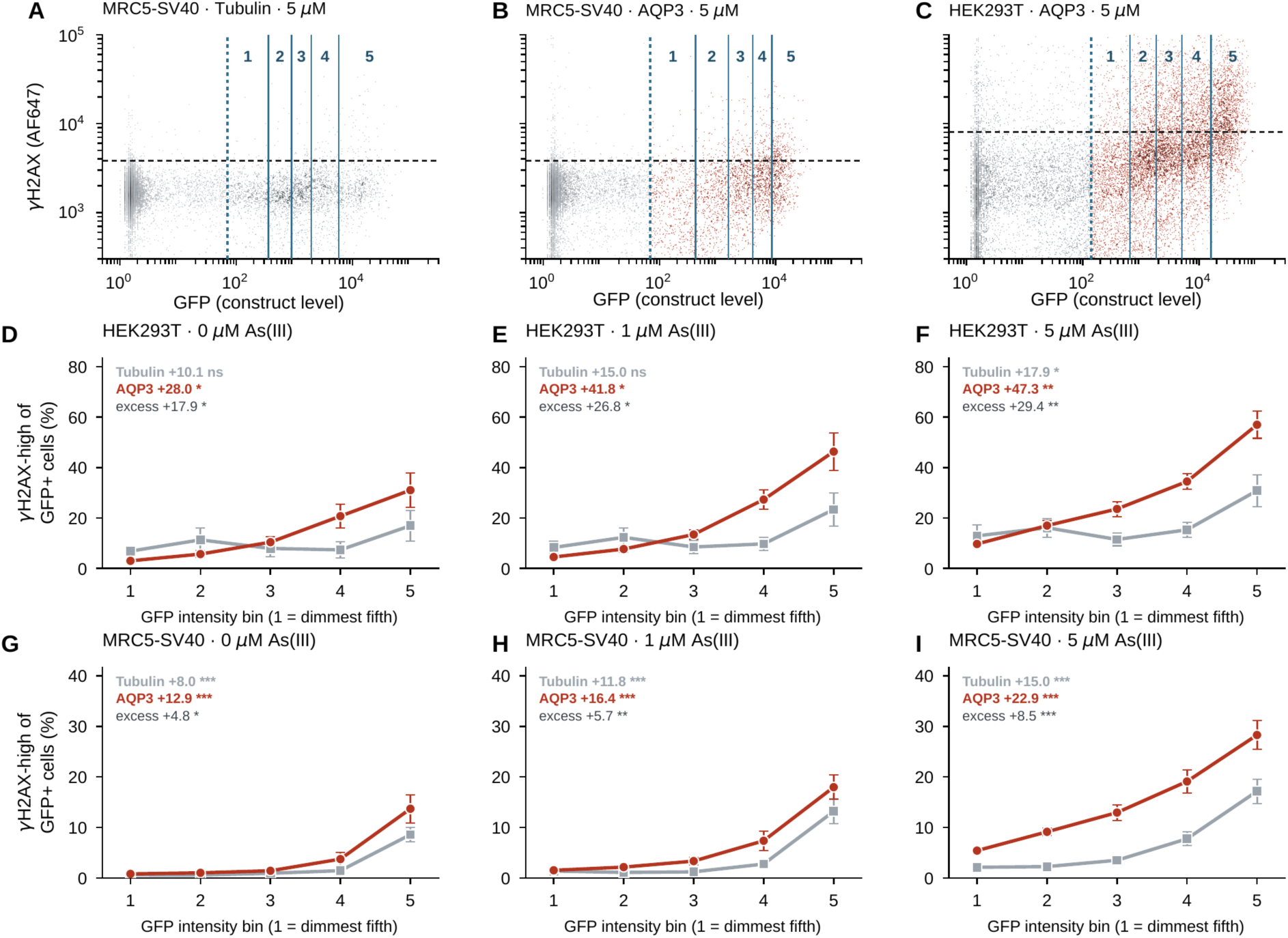
DNA damage increase with amount of AQP overproduction. Within one tube every transfected cell carries a different amount of construct, read out as GFP intensity. Splitting the GFP-positive cells of a tube into five equal-count intensity bins turns that into a dose-response in expression level measured inside a single well: brighter and dimmer cells sat in the same tube, saw the same arsenite and the same stain, so there is no plate-to-plate normalization and no transfection-efficiency confound. (A–C) How the bins are defined. GFP against γH2AX in single cells of one tube, one dot per cell, shaded by local dot density; grey, GFP-negative cells, colored, GFP-positive cells. The GFP-positive cells are cut at the quintile boundaries of their own GFP distribution into bins 1–5 (bin 1 = dimmest fifth). Dotted vertical line, the lower edge of the GFP-positive gate; solid blue lines, the four bin edges; bold numerals, bins 1–5; dashed horizontal line, that plate’s damage threshold. Both axes are logarithmic. (D–F) HEK293T and (G–I) MRC5-SV40, one panel per arsenite concentration. Grey squares, GFP-Tubulin; red circles, GFP-AQP3; points, mean ± SEM across acquisition dates with replicate tubes averaged within a date (n = 3 dates in HEK293T, 8–10 in MRC5-SV40). The y scale is shared across the three panels of a row. Labels give the bin 1 → bin 5 rise in each construct and the AQP3-minus-Tubulin excess, both paired within acquisition date and tested one-sided. Damage rises with construct level for AQP3 at every concentration, and the AQP3 gradient is steeper than the GFP-Tubulin gradient throughout: the excess grows from +17.9 to +29.4 pp across 0, 1 and 5 μM in HEK293T (P = 0.013, 0.010, 0.0016) and from +4.8 to +8.5 pp in MRC5-SV40 (P = 0.034, 0.0037, 0.0008). Two features of the control matter for interpretation. GFP-Tubulin carries a gradient of its own (+8.0 to +17.9 pp), so brightness is not inert, because more plasmid means more stress whatever it encodes, which is why the AQP3-minus-Tubulin excess rather than the raw AQP3 gradient is the channel-attributable quantity. And that excess is already present without arsenite (+17.9 pp in HEK293T, +4.8 pp in MRC5-SV40), so it is not purely arsenite-dependent; what arsenite does is make it larger, by 1.6-fold in HEK293T and 1.8-fold in MRC5-SV40 from 0 to 5 μM. Bins with fewer than 100 cells were dropped, and none were. Damage threshold, the top 0.5% of the mock-transfected distribution of that acquisition.

### N-acetylcysteine removes the AQP3-attributable excess

Arsenite generates reactive oxygen species, and scavenger rescue is the classical test of that mechanism (53). We therefore applied N-acetylcysteine (NAC, 6 mM) concurrently with arsenite in AQP3-expressing MRC5-SV40 cells, on the three acquisition plates carrying both arms (Figure 5A, B). NAC removed 47% of the GFP-AQP3 signal at 5 μM (9.04% to 5.35%; −3.69 percentage points, 90% CI −1.07 to −6.30; P = 0.027) and did nothing without arsenite (P = 0.898), while in the GFP-Tubulin control it reached significance at no dose (P = 0.735, 0.528, 0.068) (Figure 5C, D). Applied to the AQP3-minus-Tubulin excess, the quantity the channel is responsible for, NAC removed 55% at 1 μM (1.53 to 0.68 points; −0.84 points, P = 0.033) and 36% at 5 μM (5.39 to 3.47 points; −1.93 points, P = 0.034), and nothing at 0 μM, where the excess is itself near zero (Figure 5E). As a fraction of the GFP-AQP3 signal, NAC removed 43 ± 3% at 5 μM and 24 ± 9% at 1 μM, while at 0 μM the signal was if anything slightly higher with NAC (−17 ± 7%) (Figure 5F). The shape of the rescue rather than any single contrast within it carries the argument: an antioxidant acting non-specifically on the assay would suppress at every dose, including zero. Re-scoring the excess contrast at the other three damage thresholds gives the same direction throughout (at 5 μM P = 0.034, 0.042, 0.050 and 0.107; at 1 μM P = 0.033, 0.062, 0.092 and 0.121). Pairing is within acquisition plate rather than within date.

**Figure 5.**
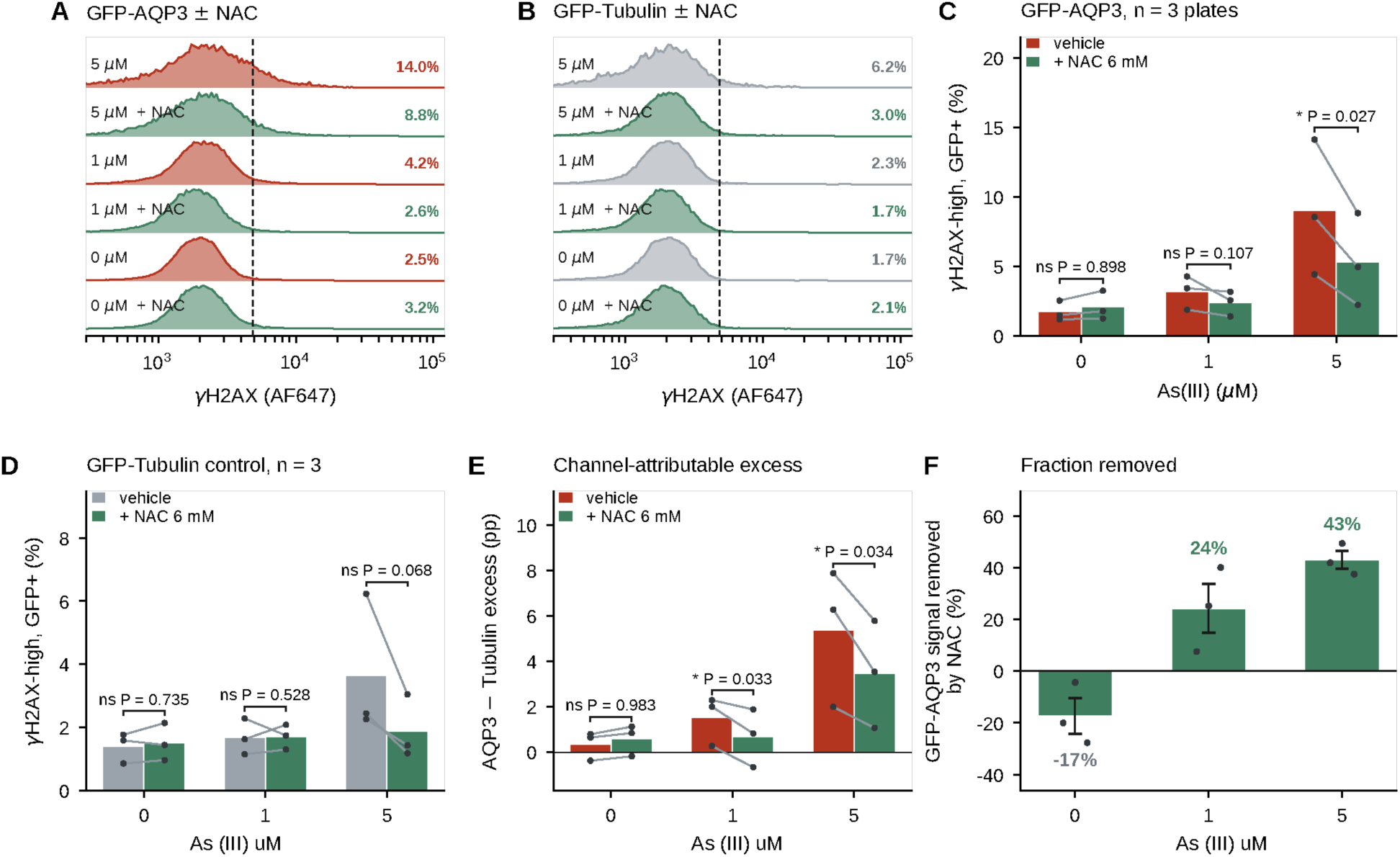
N-acetylcysteine removes the AQP3-attributable excess. (A, B) Representative γH2AX distributions in GFP-positive cells with and without 6 mM NAC at 0, 1 and 5 μM As(III), for GFP-AQP3 (A) and the GFP-Tubulin control (B) Replicate tubes of a condition are averaged as densities; dashed line, that plate’s damage threshold (top 0.5% of the mock-transfected distribution of that acquisition); the percentage above it is given at the right of each trace. (C, D) Paired vehicle/NAC values on the three acquisition plates that carry both arms, for GFP-AQP3 (C) and GFP-Tubulin (D); points and connecting lines, individual plates with replicate tubes averaged within a plate; bars, their means. NAC removes 41% of the GFP-AQP3 signal at 5 μM (9.04 → 5.35%, −3.69 pp, 90% CI −1.07 to −6.30, P = 0.027) and does nothing at 0 μM (P = 0.898). In the GFP-Tubulin control it reaches significance at no dose (P = 0.735, 0.528, 0.068). (E) The same contrast applied to the AQP3-minus-Tubulin excess, the quantity the channel is responsible for. NAC removes 55% of the excess at 1 μM (1.53 → 0.68 pp, −0.84 pp, P = 0.033) and 36% at 5 μM (5.39 → 3.47 pp, −1.93 pp, P = 0.034), and nothing at 0 μM, where the excess is itself near zero (0.36 pp) and the 0.24 pp difference is not interpretable at this n. (F) The same result as a fraction: the percentage of the GFP-AQP3 signal that NAC removes at each dose. Bars, the mean of the three per-plate percentages; whiskers, SEM; points, the individual plates. NAC removes 43 ± 3% at 5 μM and 24 ± 9% at 1 μM, and at 0 μM the signal is if anything slightly higher with NAC (−17 ± 7%). No significance marks are shown on this panel: with three plates only the 5 μM value is well determined, and the tests that support the claim are the paired contrasts in C–E. Re-scoring the excess contrast at the other three damage thresholds gives the same direction throughout: at 5 μM P = 0.034, 0.042, 0.050 and 0.107 across the 0.5, 1, 2 and 5% cutoffs, and at 1 μM P = 0.033, 0.062, 0.092 and 0.121. Pairing is within acquisition plate, not within date: two of these dates also carry a separate non-NAC plate whose tubes ran higher, and averaging those into the vehicle arm while the NAC arm came from the NAC plate alone would overstate the suppression. Only runs carrying both arms enter the analysis. All contrasts are paired and tested one-sided in the pre-specified direction (NAC lowers γH2AX), as elsewhere in this paper; two-sided P values are exactly twice those quoted. Confidence intervals are two-sided 90%, so interval and one-sided P agree. n = 3 acquisition plates.

This result also bounds a second possible route. AQP3 transports hydrogen peroxide (21) and could in principle raise oxidative DNA damage with no arsenite present. Without arsenite it did so only as a trend in MRC5-SV40 (+1.65 points, P = 0.066) though measurably in HEK293T (+4.09 points, P = 0.026), and NAC removed nothing in the absence of arsenite in either line. Constitutive peroxide influx is therefore not a sufficient explanation of the AQP3 effect in lung fibroblasts, which requires arsenic to be present, although it remains a candidate for the arsenite-independent component that HEK293T carries. The AQP3 inhibitor DFP00173 (47) at 25 μM did not reduce arsenite-associated damage in AQP3-expressing cells: over eight vehicle-versus-inhibitor pairs from three acquisition runs, paired within construct and within run, the change was +0.21 percentage points (95% CI −1.00 to +1.42; two-sided paired t-test P = 0.70; Supplementary Figure S1). Because DFP00173 and H53F may disrupt different aspects of AQP3 function, the absence of an inhibitor effect and the loss of effect in the mutant are not in conflict.

### Arsenite alone does not raise somatic mutation frequency

γH2AX reports breaks present at the moment of measurement rather than mutations fixed into the genome. To ask whether the interaction reaches the level of permanent sequence change we used duplex sequencing, which reads both strands of each original DNA duplex and drops the error floor far below that of conventional short-read sequencing (42, 43, 54). Twenty-six libraries were sequenced from MRC5-SV40 cells across two capture panels: 19 on a custom ∼200 kb panel tiling the EGFR locus and 7 on the TwinStrand human-muta-v1.0 48 kb mutagenesis panel.

The method separates a real mutation from an amplification or sequencing error by requiring both strands of the original duplex to agree adapters carrying a strand-distinguishing dual unique molecular identifier are ligated to each fragment, reads are grouped by that identifier, a consensus is called for each strand, and only changes present in both single-strand consensuses are retained (Figure 6A). Two controls bound what the assay reports. Positions called in three or more independent libraries cannot be independent de novo events: 289 such positions occur where an independence null predicts about one, and removing them discards about a third of all somatic calls. And the six-channel spectrum is indistinguishable between arsenite-treated and untreated AQP3-positive cells (cosine similarity 0.9967), so nothing in what follows rests on a spectral shift.

**Figure 6.**
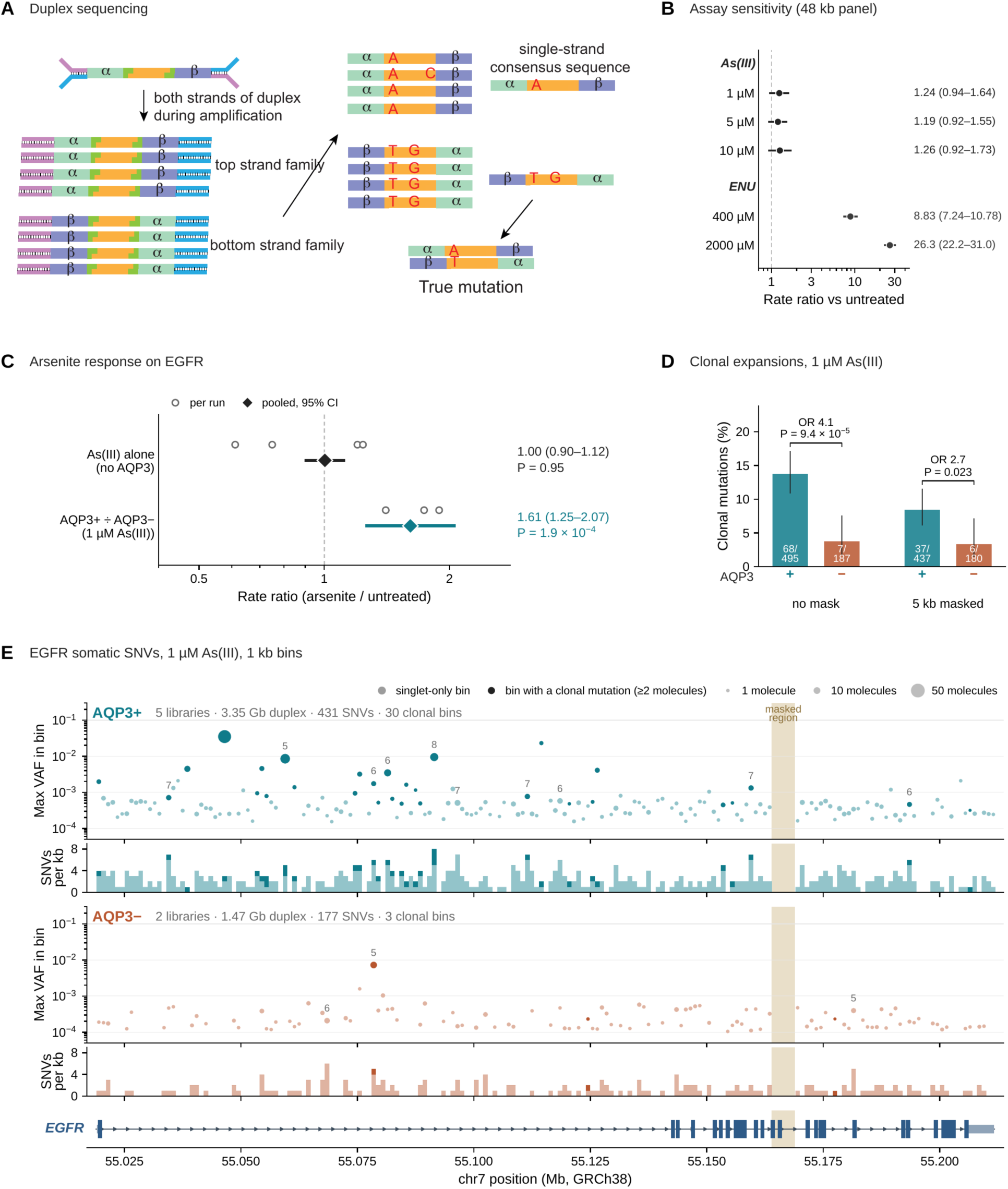
Duplex sequencing: principle, sensitivity and the AQP3-dependent arsenite response at EGFR. (A) A fragment is ligated to adapters carrying a strand-distinguishing dual unique molecular identifier and amplified; reads are grouped by that identifier and a consensus called for each strand, so a change present on both single-strand consensuses is scored as a true mutation and one present on a single strand is discarded as an amplification or sequencing error. (B) Assay sensitivity on the 48 kb TwinStrand human-muta-v1.0 panel in parental cells: rate ratio against untreated for arsenite at 1, 5 and 10 μM and for N-ethyl-N-nitrosourea at 400 and 2000 μM, with 95% confidence intervals. (C) Arsenite response on the EGFR panel: arsenite alone (1 versus 0 μM) in cells without AQP3 overexpression, and the genotype-dependent response (AQP3+ ÷ AQP3− at 1 μM As(III)); open circles, individual sequencing runs; diamond, pooled estimate with 95% confidence interval, by generalized least squares for the genotype comparison because its contrasts share a denominator; dashed line, rate ratio 1. (D) Share of somatic calls that are clonal, defined as supported by two or more duplex molecules, at 1 μM As(III) by AQP3 genotype, before and after masking the 5 kb misalignment window; error bars, Wilson 95% confidence intervals; fractions inside the bars give clonal over total calls. (E) Somatic single-nucleotide variants at 1 μM As(III) across the EGFR target in 1 kb bins, AQP3-positive above and AQP3-negative below; upper track, maximum variant allele fraction in each bin with point area scaled to supporting molecules and saturated points marking bins carrying a clonal mutation and pale points bins of singlets only; lower track, variants per kilobase; shaded column, the masked 5 kb region; gene model, EGFR (GRCh38). Filters throughout: somatic calls only (VAF < 0.01), VarDict PMEAN ≥ 8, recurrent-position blacklist and per-read-length mappability mask; log rate ratios pooled by fixed-effect inverse-variance weighting, or by generalized least squares where contrasts share a denominator; P values two-sided.

The assay resolves a mutagen when one is present. On the 48 kb panel, ENU raised mutation frequency 8.83-fold at 400 μM (95% CI 7.24-10.78) and 26.3-fold at 2000 μM (22.2-31.0) over untreated parental controls, monotonically in dose (Figure 6B). Against that, arsenite is flat: 1.24-fold at 1 μM, 1.19 at 5 μM and 1.26 at 10 μM, none reaching significance.

Four independent contrasts of 1 μM against 0 μM in cells that do not overexpress an aquaglyceroporin gave rate ratios of 1.20, 0.61, 0.75 and 1.24, pooling to 1.00 (0.90-1.12; P = 0.95) with I^2^ = 85% (Figure 6C, upper row). The heterogeneity is the point rather than a nuisance: the two unsorted parental contrasts lie above one and the two sorted GFP-negative contrasts below it, so the pooled value states that the four contrasts do not agree on a direction, not that each is null. No contrast in the set approaches the magnitude the same assay resolves for ENU. Arsenite alone therefore produced no consistent change in mutation frequency at any dose to 10 μM, which bounds any effect rather than demonstrating a null.

This is what the mechanistic literature predicts. Arsenite is mutagenic at the thymidine kinase locus only at high doses yet enhances ultraviolet mutagenesis more than additively by inhibiting repair of pyrimidine dimers(40); arsenic alone did not raise mutation burden in mouse skin while more than doubling the ultraviolet mutational burden (41); and arsenite binds and inhibits zinc-finger repair proteins (39) and lowers XPC protein in human lung fibroblasts (55). A dissociation between a damage signal and a mutational one is expected, and it is the reason a transporter that raises internal dose, rather than arsenite itself, is the variable worth testing.

### The mutational response to arsenite differs by AQP3 genotype

We then normalized each genotype to its own untreated control from the same sequencing run and compared genotypes, a difference-in-differences that removes run, read length and capture batch from the contrast (Figure 6C, lower row). The three within-run comparisons gave 1.89 (1.14-3.13; P = 0.014), 1.41 (1.01-1.96; P = 0.043) and 1.74 (1.19-2.53; P = 0.0040), pooling to 1.61 (1.25-2.07; P = 1.9 × 10-4) by generalized least squares, which accounts for the two comparisons sharing one AQP3-negative denominator. The estimate is stable under masking: 1.61 with the per-read-length mappability mask alone, 1.57 with a 1.5 kb hypermutated window additionally excluded, and 1.55 with the wider 5 kb misalignment region excluded.

Two properties qualify that number, and we state them with it. First, the magnitude and sign depend on which cells serve as the reference. Against GFP-negative cells sorted from the same transfection, the AQP3-attributable response is 1.43-fold (1.14-1.81; P = 0.0023) unpaired and 1.61-fold paired within run; against never-transfected parental cells it is 0.86 (0.69-1.06) at 1 μM. We prefer the GFP-negative reference because it alone is matched to the AQP3-positive fraction for transfection, sorting and read length matching, not outcome, motivates the choice but the choice was made after the analysis rather than in advance, and the parental comparison is tabulated alongside it. Second, the 2 × 76 bp unit, which gives the largest of the three within-run values, is the one that does not survive the region masks (1.52, P = 0.13 with the 1.5 kb window excluded), so the stability of the pooled estimate under masking is a property of the pool and not of every unit in it.

A recurrent-position blacklist removes somatic calls appearing at the same position in three or more independent libraries, which cannot be independent de novo events: 289 such positions across the 19 EGFR libraries at PMEAN ≥ 8, where chance predicts about one, together accounting for roughly a third of somatic calls. A regional scan of the EGFR target identified a 5 kb window, chr7:55 163 933-55 168 933, in which T>A accounts for 27.5% of calls against 5.0% elsewhere, 117 of 142 positions are singletons, and which is hypermutated in untreated control libraries as well; that combination identifies reads mismapping from a paralogous sequence rather than mutagenesis. This one window accounted for an apparent 14% arsenite elevation in an earlier unpaired analysis, which fell from 1.14 to 1.05 once the window was excluded.

Four further features limit the genotype comparison. The intervals assign Poisson variance to each contrast, treating every interrogated duplex base as an independent trial, whereas the library is the correct replicate unit; on the unpaired five-against-five comparison a library-level rank test gives P = 0.42 where the Poisson test gives 0.038. Two of the three comparisons in Figure 6C share the same AQP3-negative denominator samples and are not three independent biological replicates; the pooled estimate therefore models that shared denominator by generalized least squares, and because those two comparisons are also two DNA preparations of a single sorted cell population rather than two biological replicates, even that understates the correlation. The heterogeneity in Figure 6C splits by sorting status rather than at random. And although the GFP-positive and GFP-negative fractions are matched for everything we imposed, they differ in whether the cell took up and expressed plasmid at all rather than only in AQP3. No signature analysis is reported at these mutation counts, and the six-channel spectrum shows no difference between arsenite-treated and untreated AQP3-positive cells (cosine similarity 0.9967). We therefore present these duplex data as supporting and provisional.

Because the panel carrying this comparison tiles EGFR across 192 780 bp of chr7:55 018 933-55 211 713, the AQP3-dependent increment is measured at EGFR. It is a burden across the locus and not a driver signal: exons are 9793 bp, 5.1% of the target, and are not enriched for somatic sites in any arm, with no clustering at the kinase domain or at any known EGFR driver position. At these mutation counts the design is powered for burden and not for hotspots, so the absence of driver-position clustering is uninformative. Whether the increment is specific to EGFR cannot be answered here, because the second panel carries no AQP3 arm.

Two further features of the same libraries bear on how that increment arises. Clonal calls are more common in AQP3-positive than in AQP3-negative libraries under arsenite: at 1 μM As(III), 13.7% of somatic calls in AQP3-positive libraries were clonal (68 of 495) against 3.7% in AQP3-negative libraries (7 of 187), an odds ratio of 4.1 (P = 9.4 × 10⁻⁵), and the difference survives masking the 5 kb misalignment window, where the shares are 8.5% (37 of 437) and 3.3% (6 of 180), an odds ratio of 2.7 (P = 0.023) (Figure 6D). A clonal call reports one ancestral event sampled repeatedly rather than several independent ones, so this is a caution as much as a result: part of the raw burden difference between genotypes reflects the expansion of a smaller number of founding mutations, which is why the frequency estimate above is reported only after the recurrent-position blacklist and the mappability mask. Along the target itself the variants are spread across the locus in both genotypes rather than concentrated at any position: 431 somatic single-nucleotide variants in five AQP3-positive libraries over 3.35 Gb of interrogated duplex territory, of which 30 one-kilobase bins carry a clonal mutation, against 177 variants in two AQP3-negative libraries over 1.47 Gb with three such bins (Figure 6E).

### Arsenite induces a graded oxidative-stress program, and the response is unevenly distributed across cells

We applied SHERRY (51) to pools of ten sorted MRC5-SV40 cells in a two-by-two design (AQP3-positive and GFP-negative, 0 and 1 μM As (III)) together with a dose ladder acquired within a single sequencing run (Figure 7A). Pool size sets what the design can resolve. Ten cells is a deliberate compromise: a minority response stays visible (one responding cell in ten lifts pooled HMOX1 about 1.6-fold) but no pool can reach the fully responding level. Exonic read fraction was lower in AQP3-positive pools than in GFP-negative ones (medians 13.9% without arsenite and 15.3% at 1 μM, against 20.5%, 26.8% and 34.6% at 0, 1 and 10 μM), and within the GFP-negative arm it rose with dose, so every gene-level fit below is adjusted for sequencing run; whether this reflects genuine intron retention would require isoform-level quantification we have not performed.

**Figure 7.**
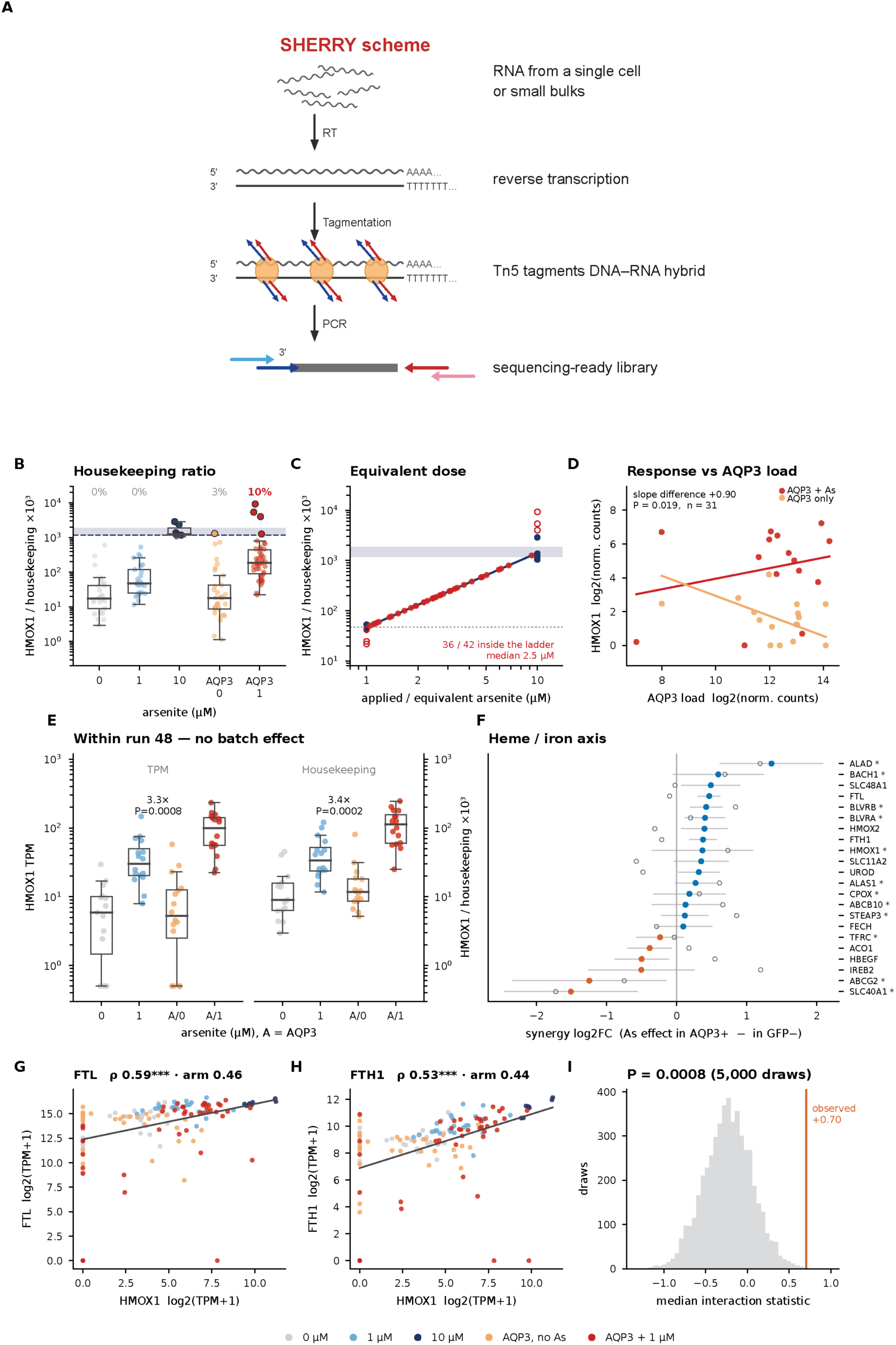
Low-input RNA sequencing resolves a graded oxidative-stress response and its dependence on AQP3 load. (A) SHERRY workflow. RNA from a single cell or a small pool is reverse-transcribed with oligo-dT; Tn5 tagments the resulting RNA/DNA hybrid directly, so no second-strand synthesis is needed, and the product is amplified into a sequencing-ready library in one tube. (B) HMOX1 normalized to housekeeping transcripts across the arsenite dose ladder and the two AQP3 arms; box, interquartile range; points, individual libraries; shaded band, the interquartile range of the seven 10 μM libraries and dashed line its lower edge; percentages, the share of each group reaching that band; all 135 ten-cell libraries. (C) Equivalent arsenite dose: each AQP3 + 1 μM library’s housekeeping ratio read against a log-log fit to the run-220 libraries at 1 and 10 μM (slope 1.48 decades per decade; the 5 μM rung is excluded); open circles, libraries falling off the ladder, drawn at the rung they passed; an x position is inferred rather than an applied dose; 36 of 42 libraries fall inside the ladder, with a median equivalent of 2.5 μM. (D) HMOX1 against AQP3 load across libraries, arsenite-treated and untreated; lines, fitted slopes, with the difference in slope and its P value. (E) HMOX1 within sequencing run 48 alone, as TPM and as the housekeeping ratio, across untreated, 1 μM, AQP3 without arsenite and AQP3 with 1 μM; fold changes and P values above the brackets. (F) Heme and iron-handling genes ranked by the synergy log2 fold change, the arsenite effect in AQP3-positive pools minus that in GFP-negative pools; bars, ±1 standard error; filled points, the within-run estimate; open points, the batch-adjusted estimate across 76 libraries; asterisk, genes of the same sign in both. (G, H) FTL and FTH1 against HMOX1 across all libraries, colored by condition; ρ, partial Spearman correlation with HMOX1 across all 135 libraries, adjusted for sequencing depth, genes detected and run; arm, the same correlation inside the AQP3 + 1 μM arm alone. (I) Permutation null for the median interaction statistic of the heme/iron set against 5,000 expression-matched random sets; line, the observed value.

Arsenite produced a canonical, dose-graded NRF2 response, and expressing HMOX1 against a housekeeping denominator makes that ladder comparable across libraries of unequal depth. The ratio rises monotonically from 0 to 1 to 10 μM applied arsenite, and the fraction of libraries sitting above the saturation band rises with it, from none at 0 and 1 μM to 3% with AQP3 alone and 10% with AQP3 and 1 μM arsenite together (Figure 7B). Transcription therefore corroborates the oxidant response invoked in the N-acetylcysteine experiment. Reading each library back against that internal ladder converts its HMOX1 level into an equivalent applied dose: 36 of 42 libraries fall inside the ladder, with a median equivalent dose of 2.5 μM in cells that received 1 μM (Figure 7C). The response scales with how much channel a pool carries. HMOX1 rises with AQP3 load in arsenite-treated pools and falls with it in pools given AQP3 alone, a slope difference of +0.90 (P = 0.019, n = 31; Figure 7D). Within the single sequencing run that contains all four groups of the design, where batch is removed entirely, AQP3 raised HMOX1 3.3-fold at 1 μM on TPM (P = 0.0008) and 3.4-fold on the housekeeping ratio (P = 0.0002), while AQP3 without arsenite was indistinguishable from untreated (Figure 7E). That shape is what sensitization predicts rather than a constitutive effect.

What the pooled design does show is over-dispersion rather than a uniform shift. AQP3-plus-arsenite pools sit mostly inside the single-condition range with a few far outside it, and variance was inflated on an oxidative DNA repair signature (Brown-Forsythe P < 0.001; 4 of 19 pools above the 90th percentile of the single conditions) and on an integrated stress signature (P = 0.010; 2 of 19), with the NRF2 core signature in the same direction but not significant (P = 0.083). This is the distributional signature of a mixed responder and non-responder population. One gene behaves as a combined insult predicts. Of 2,905 testable genes, SQSTM1 alone is significantly raised above both single conditions: +0.74 log2 over arsenite alone (adjusted P = 0.024) and +1.21 log2 over AQP3 alone (adjusted P = 1.6 × 10⁻⁷). SQSTM1 encodes p62, which sequesters KEAP1 and thereby stabilizes NRF2, placing the one gene that exceeds both single conditions on the pathway the N-acetylcysteine experiment implicates. With a single survivor, this is a lead rather than a demonstrated interaction. At gene level the design cannot localize the effect. No gene reached a 5% false discovery rate for an AQP3-by-arsenite term (smallest adjusted P = 0.23 across 120 measurable panel genes), and the panel was not enriched over expression-matched random sets (P = 0.24 on hit count; P = 0.066 on median interaction statistic). Twelve genes reach nominal significance; only TFRC survives dropping any single pool (interaction +0.80, P = 0.0078; worst leave-one-out refit P = 0.037), where HMOX1 does not (+0.97, P = 0.0076; worst refit P = 0.074).

A wider panel of heme and iron-handling genes places the interaction on that axis. Ranking the heme and iron genes by their synergy log2 fold change, the arsenite effect in AQP3-positive pools minus the same effect in GFP-negative pools, puts the heme-synthesis and heme-degradation arm at the positive end, with ALAD, BACH1, BLVRB, BLVRA, HMOX1, ALAS1, CPOX, ABCB10 and STEAP3 above zero and of the same sign in the batch-adjusted estimate, and the iron-export arm at the negative end, with SLC40A1 and ABCG2 below it (Figure 7F). No individual gene in the set survives correction for multiple testing, so the set-level test below rather than any single gene carries this claim. HMOX1 does not move alone: across libraries its level tracks both ferritin subunits, ρ = 0.59 for FTL and ρ = 0.53 for FTH1, and the association persists within treatment arm at ρ = 0.46 and 0.44, so it is not produced by the dose contrast alone (Figure 7G, H). Tested as a set against 5,000 expression-matched random gene sets, the observed median interaction statistic of +0.70 falls outside the permutation distribution (P = 0.0008; Figure 7I). The effect of AQP3 on the arsenite response is therefore not confined to a single NRF2 target but runs across the heme-degradation and iron-handling program that HMOX1 belongs to. These remain leads for a powered design rather than established interactions.

### The response is shared with AQP9 and AQP10 and depends on the host cell

Several aquaglyceroporins conduct trivalent arsenic (16, 18, 19)so we tested AQP7, AQP9 and AQP10 against the GFP-Tubulin control acquired on the same plate, in all three lines and across the full 0, 1 and 5 μM series (Figures 8 and 9). In MRC5-SV40, AQP3 was raised at every dose (+1.51, +2.58 and +7.69 percentage points; P = 0.047, 0.013 and < 0.0001, n = 10-11 plates). These contrasts are paired within acquisition plate rather than within date, so the AQP3 values differ slightly from the date-paired estimates of Figure 1. AQP7 and AQP9 were raised without arsenite (+2.86 points, P = 0.029, and +2.74 points, P = 0.071) and rose further with it, whereas AQP10 sat if anything below the control at 0 and 1 μM (−1.03 and −1.14 points) and then rose furthest at 5 μM (+13.4 points). Spontaneous and arsenite-driven activity are separable properties of a paralogue rather than one property (Figure 8G). In HEK293T, AQP3 was again raised at every dose, by the amounts already given above, while AQP9 and AQP10 were raised only once arsenite was present (AQP9 +5.41 and +5.36 points at 1 and 5 μM, P = 0.014 and 0.030; AQP10 +5.67 and +5.29 points, P = 0.060 and 0.017) and AQP7 at no dose (−1.44, +0.87 and −1.13 points) (Figure 8H). The three MRC5-SV40 paralogue bars at 5 μM (AQP7 +6.15, AQP9 +11.80, AQP10 +13.44 points) rest on two plates each and carry no test; they are shown because the two plates agree closely, not as evidence in themselves. In RKO, AQP9 raised the γH2AX-high fraction by 9.05 points (P = 0.0015) and AQP10 by 7.59 points (P = 0.0002) at 5 μM, while AQP7 did not (Figure 9D).

**Figure 8.**
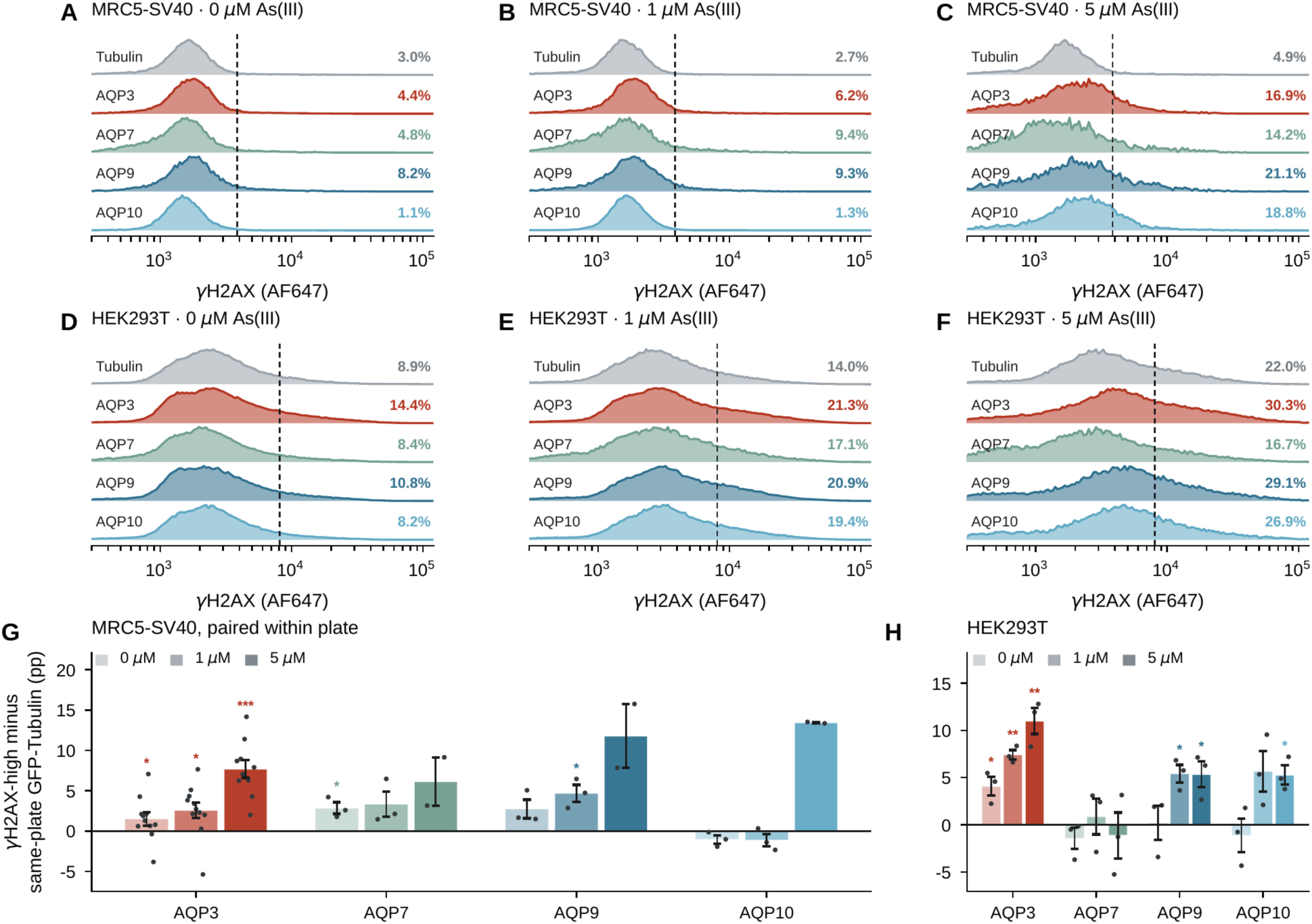
The aquaglyceroporin paralogues in MRC5-SV40 and HEK293T. (A–C) Representative γH2AX distributions in GFP-positive MRC5-SV40 cells carrying GFP-Tubulin or GFP-AQP3, -AQP7, - AQP9 or -AQP10, at 0, 1 and 5 μM As(III). (D–F) The same in HEK293T (13 October 2023 plate). Replicate tubes of a condition are averaged as densities; dashed line, that plate’s damage threshold (the top 0.5% of the mock-transfected, antibody-stained single-cell distribution of that acquisition); the percentage above it is given at the right of each trace. (G, H) Each construct minus the GFP-Tubulin control acquired on the same plate, one point per acquisition plate; bars, their means; whiskers, SEM across plates. Stars mark a significant one-sided paired test; bars without a star did not reach significance, and the n behind each is given below. In MRC5-SV40 AQP3 is raised at every dose (+1.51, +2.58, +7.69 pp; P = 0.047, 0.013, < 0.0001, n = 10–11 plates). AQP7 and AQP9 are raised without arsenite (+2.86 pp, P = 0.029 and +2.74 pp, P = 0.071) and rise further with it, whereas AQP10 is if anything below the control at 0 and 1 μM (−1.03 and −1.14 pp) and then rises furthest at 5 μM (+13.4 pp): spontaneous and arsenite-driven activity are separable properties of a paralogue, not one property. In HEK293T AQP3 is again raised at every dose (+4.09, +7.41, +11.00 pp; P = 0.026, 0.0023, 0.0078), AQP9 and AQP10 only once arsenite is present (AQP9 +5.41 and +5.36 pp at 1 and 5 μM, P = 0.014 and 0.030; AQP10 +5.67 and +5.29 pp, P = 0.060 and 0.017), and AQP7 at no dose (−1.44, +0.87, −1.13 pp). Pairing is within acquisition plate, not within date. Two of the MRC5-SV40 dates also carry a separate NAC plate, and pairing by date allowed that plate’s GFP-Tubulin tubes into the control arm while the paralogues sat only on the paralogue plate. Within-plate pairing removes that at the cost of n. The n behind each bar in G is: GFP-AQP3, 11 plates at 0 and 1 μM and 10 at 5 μM; AQP7, AQP9 and AQP10, three plates at 0 and 1 μM and two at 5 μM. In H every contrast rests on three plates. The three MRC5-SV40 paralogue bars at 5 μM (AQP7 +6.15, AQP9 +11.80, AQP10 +13.44 pp). Tests are one-sided paired t-tests of the pre-specified direction (an aquaglyceroporin raises γH2AX); two-sided P values are exactly twice those quoted. Where a plate carries two tubes per arm, GFP-AQP3 in every line and GFP-AQP10 in MRC5-SV40, those tubes are averaged within the plate before testing. As a secondary analysis, a linear mixed model with acquisition plate as a random intercept uses both tubes without treating them as independent replicates, tested on degrees of freedom. The point estimates are unchanged and the intervals tighten slightly: MRC5-SV40 AQP3 P = 0.021, 0.0028 and < 0.0001 at 0, 1 and 5 μM (against 0.047, 0.013 and < 0.0001), and HEK293T AQP3 P = 0.021, 0.0062 and 0.0081 (against 0.026, 0.0023 and 0.0078). For every contrast with a single tube per arm per plate, all of AQP7 and AQP9 and AQP10 outside MRC5-SV40, the mixed model is algebraically identical to the paired test shown.

**Figure 9.**
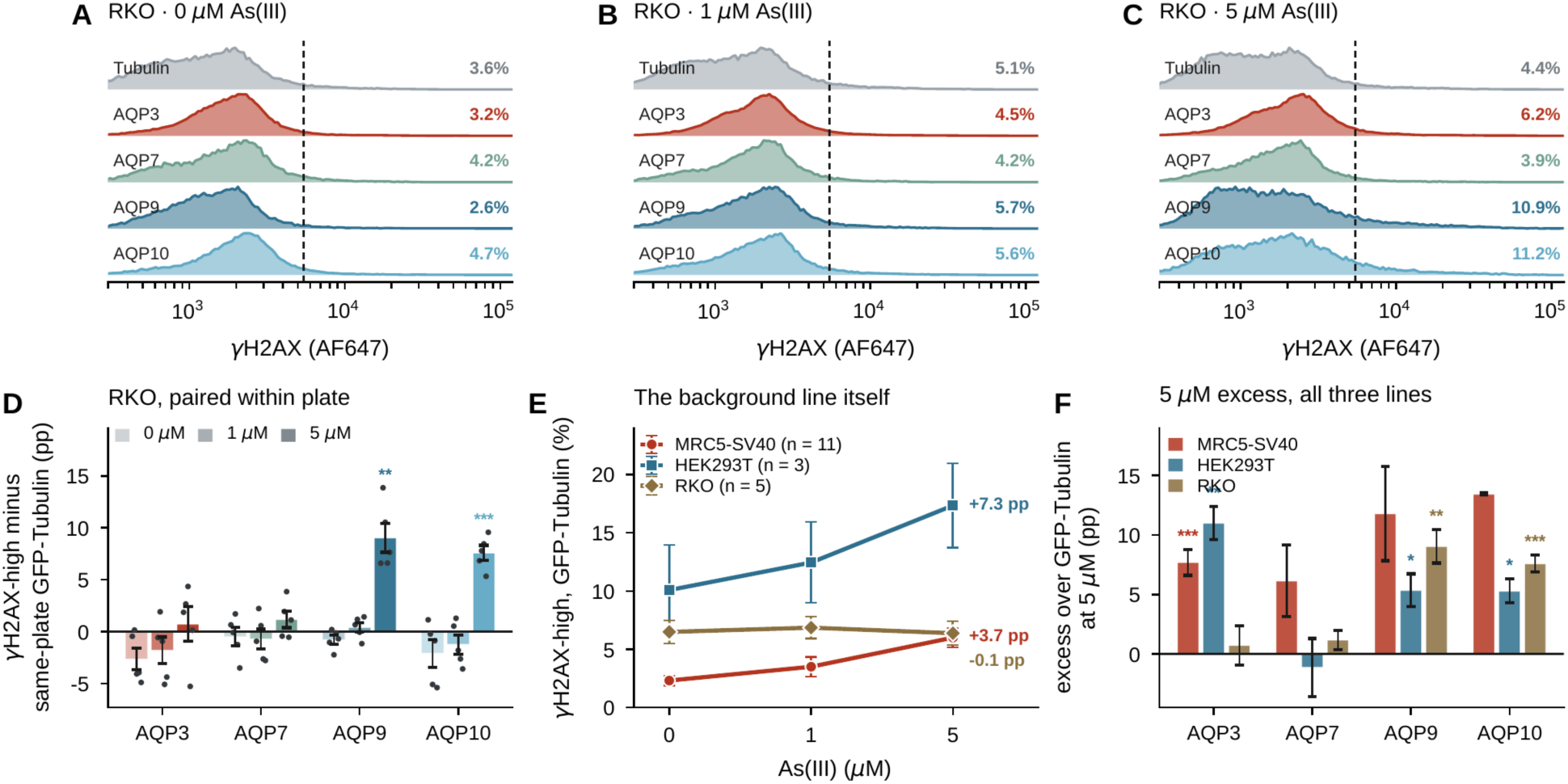
RKO separates AQP3 from the other paralogues. (A–C) Representative γH2AX distributions in GFP-positive RKO cells at 0, 1 and 5 μM As(III), plotted as in Figure 8. Of the five RKO plates this one sits closest to the pooled result in D–F, by the summed absolute deviation of its GFP-Tubulin dose response and its four 5 μM excesses from the five-plate means; it also carries the largest GFP-positive populations (18 000–53 000 cells per tube). Individual plates differ: the GFP-Tubulin arm changes by −3.3 to +3.4 pp from 0 to 5 μM across the five, which is the spread behind the flat pooled line in E, so no single plate should be read as the effect. (D) Each construct minus the GFP-Tubulin control acquired on the same plate, across the five RKO acquisition plates; bars, means; whiskers, SEM; points, the individual plates. Stars mark a significant one-sided paired test; unmarked bars did not reach it. Every RKO contrast rests on five plates. AQP3 confers no excess at any dose (−2.65, −1.82, +0.74 pp; P = 0.97, 0.88, 0.34), and neither does AQP7. AQP9 and AQP10 do, and only with arsenite (+9.05 pp, P = 0.0015 and +7.59 pp, P = 0.0002 at 5 μM). (E) The GFP-Tubulin background itself across dose in the three lines. It rises with arsenite in MRC5-SV40 (+3.7 pp from 0 to 5 μM, P = 0.0001, n = 11 plates) and in HEK293T (+7.3 pp, n = 3), and does not move at all in RKO (−0.1 pp, P = 0.93, n = 5). RKO is therefore not a less arsenite-sensitive version of the other two lines with everything else held constant: the untransfected background does not respond, yet AQP9 and AQP10 still produce a large arsenite-dependent excess in it. (F) The 5 μM excess over GFP-Tubulin, construct by construct, in all three lines; error bars, SEM across plates; stars mark a significant one-sided paired test. The three MRC5-SV40 paralogue bars rest on two plates and carry no test (see Figure 8). AQP9 and AQP10 are active in all three; AQP3 is active in MRC5-SV40 and HEK293T and not in RKO; AQP7 is active in neither HEK293T nor RKO and rests on two plates in MRC5-SV40. Conventions and tests as in Figure 8, including the mixed-model secondary analysis: in RKO the only contrast with two tubes per arm is GFP-AQP3, whose P values become 0.99, 0.95 and 0.26 at 0, 1 and 5 μM (against 0.97, 0.88 and 0.34); it remains without effect at every dose.

The same measurement across the construct panel shows the ordering directly in the raw distributions. Without arsenite the γH2AX distributions of all five constructs overlap; at 5 μM they separate, with AQP3, AQP9 and AQP10 shifting a substantial right-hand fraction across the threshold while GFP-Tubulin does not (Figure 8A, C), and the AQP3 dose series moves progressively rightward (Figure 1A).

RKO makes the sharpest form of the observation available. The GFP-Tubulin background rises with arsenite in MRC5-SV40 (+3.7 points from 0 to 5 μM, P = 0.0001, n = 11 plates) and in HEK293T (+7.3 points, n = 3), and does not move at all in RKO (−0.1 points, P = 0.93, n = 5) (Figure 9E). RKO is therefore not a less arsenite-sensitive version of the other two lines with everything else held constant: its untransfected background does not respond, yet AQP9 and AQP10 still produce a large arsenite-dependent excess in it (Figure 9A-D, F). An aquaglyceroporin can therefore create an arsenite dose-response in a cell that has none of its own, which is a stronger statement than amplification of an existing one.

In the same line and against the same comparator, AQP3 conferred no excess at any dose (−2.65, −1.82 and +0.74 points; P = 0.97, 0.88 and 0.34, n = 5 plates) and sat below the control at the two lower doses. RKO is not an unresponsive system, because AQP9 and AQP10 were strongly active in it. Transfection efficiency in RKO was the lowest of the three lines (median 12% GFP-positive against 22% in MRC5-SV40 and 65% in HEK293T), but within a plate the AQP3 and GFP-Tubulin arms were matched (+3.1 points, n = 5, P = 0.50) and efficiency was comparable across constructs in the line, so it does not account for a result confined to AQP3; arsenite-associated cell loss was also smallest in this line, so the 5 μM cytotoxicity caveat applies least where the negative result was obtained.

The response is therefore neither unique to AQP3 nor a property of the family as a whole: AQP7, which conducts arsenite in *Xenopus* oocytes (18), produced no detectable difference in either line. AQP10, by contrast, was among the strongest, even though the same heterologous assay reported little or no As(OH)₃ permeability for the human protein (22) and the only positive evidence for arsenic handling by an AQP10 orthologue comes from zebrafish (56).

## DISCUSSION

Chronic iAs exposure affects more than 200 million people, and susceptibility at equivalent intake varies by an amount metabolism alone has not explained (3, 11, 12). We find that the γH2AX gap between AQP3-expressing and control cells widens with arsenite dose: AQP3 changes the slope of the exposure-response relationship rather than adding a constant amount of damage. The distinction matters for risk: a constitutive source of damage contributes equally at every exposure and largely cancels between exposed and unexposed people, whereas a modifier of the slope alters the genotoxic consequence of a fixed exposure, which is what exposure-response modeling estimates and what one population-average slope cannot represent. Additivity on the percentage-point scale is the null under test; of the two dose steps only the second survives it, and against a multiplicative null the two separate further, the AQP3-to-control ratio being flat from 0 to 1 μM before rising to 2.36-fold. The interaction holds on either null from 1 to 5 μM, and we do not call the first step synergy.

Our results agree with the human-cell evidence that AQP3 gates arsenite entry and disagree with the one heterologous flux measurement that addressed hAQP3 directly (22, 23). Three differences may reconcile them: oocyte flux is read over minutes at a single pH whereas our endpoint integrates 72 h in a native human membrane, and AQP3 gating is itself pH- and redox-sensitive (57); heterologous expression need not reproduce that membrane’s lipids, modifications or partners; and a damage endpoint reports the product of influx, efflux, metabolism and repair, so a modest permeability difference can produce a large downstream one. These data cannot adjudicate transport, which requires the measurement we did not make.

AQP3 is not the only aquaglyceroporin that confers sensitivity here, and the breadth cuts both ways. AQP9 and AQP10 were active in RKO cells where AQP3 was not, creating a dose-response in a line that had none of its own, the clearest evidence in this work that the transporter, not the exposure, sets the damage. That fits an AQP9 literature in which its expression sets arsenic uptake and sensitivity in leukemia cells (58) and forced expression abolishes acquired resistance in lung cancer cells (59). For AQP10 no mammalian arsenic data exist, the only comparable work being in zebrafish (56), and mouse Aqp10 is a pseudogene a plausible reason the field has not looked. Against that, the breadth costs the specific motivation for studying AQP3, its identification as a lung cancer-associated, DNA damage-promoting protein (28, 29, 32). We do not regard the MRC5-SV40 paralogue ranking as established: those rows rest on two experiments each and no construct’s expression was measured.

Duplex sequencing addressed the gap γH2AX leaves and produced two results pointing in different directions. Arsenite alone produced no consistent change in mutation frequency in cells without AQP3 overexpression, at any dose to 10 μM, in an assay that resolved 8.8-fold and 26-fold responses to ENU in the same cells; the four contrasts disagree in direction rather than agreeing on zero, so this bounds an effect rather than demonstrating a null. The genotype-dependent term is weaker than it first appears. It inverts to 0.86 against never-transfected parental cells, and that reference was chosen after the analysis; a quantity whose sign depends on a post hoc reference cannot carry a mechanistic claim. Much of an apparent effect can also be architectural: one 5 kb misalignment window, hypermutated in untreated controls too, accounted for an entire 14% arsenite elevation in an unpaired analysis, and a blacklist of 289 positions recurring in three or more libraries where independence predicts about one removed about a third of all somatic calls. Independent biological replicates in all four arms, on one panel and read length with the reference fixed in advance, would convert this into a result.

The locus is not incidental. The panel tiles EGFR, and two independent lines of evidence already connect EGFR and AQP3 to lung adenocarcinoma risk: transcriptome-wide association identifies a susceptibility locus at 9p13.3 driven by higher predicted AQP3 expression (29), and lung proteomics integrated with GWAS identifies AQP3 protein as one of two adenocarcinoma-specific susceptibility proteins (32). If a transporter that is itself a susceptibility gene converts a common exposure into a mutational one at EGFR, the two observations describe one mechanism rather than two coincidences. These data establish an AQP3-dependent burden difference within EGFR under one reference; they do not establish specificity to EGFR, and they do not show driver-position clustering, which a burden-powered study would not detect.

That suggests a testable hypothesis the field does not currently test. EGFR-mutant lung adenocarcinoma is distributed unevenly across populations (60–62), and so is inorganic arsenic exposure (63–66). An exposure that is not mutagenic alone, but becomes mutagenic where aquaglyceroporin expression is high, would produce a population-structured excess without ever appearing as a mutagen in a single-agent assay. Testing it requires EGFR mutation status, an arsenic exposure biomarker and AQP3 expression or 9p13.3 genotype measured in the same individuals. A comparison of national EGFR mutation frequencies against arsenic-province status cannot do it: such an average runs over exposure, ancestry, smoking prevalence, histological classification and screening intensity at once, neither variable is measured in the people whose tumors were sequenced, and the geographic variation in EGFR frequency already tracks East Asian ancestry markers with no evident arsenic connection.

Three observations converge on a single-cell reading, which we advance as a model rather than a demonstrated mechanism. The γH2AX endpoint is itself a count of cells above a threshold, and AQP3 roughly doubles that count at 5 μM while leaving untransfected neighbors in the same well unaffected; the ten-cell pools are over-dispersed rather than shifted; and the duplex data locate a mutational difference in the genotype carrying the channel. The model is that AQP3 raises cytoplasmic arsenite unevenly transient overexpression spans roughly a hundredfold in construct level, so a subset of cells experiences an internal dose far above the nominal one and crosses into a damaged, oxidant-stressed and mutable state. A sensitized subpopulation is the natural unit for a carcinogenesis argument, since a tumor originates in one cell rather than in a population average, and it would be invisible to any bulk assay. The evidence is convergent in direction across three assays and statistically independent in none.

The shape of the N-acetylcysteine rescue, rather than any contrast within it, is the strongest mechanistic evidence here: nothing without arsenite, partial suppression at 1 μM, half the damage removed at 5 μM. The pattern fits the oxyradical production arsenite drives (53) without identifying the species, and two routes remain open. Cytoplasmic arsenite generates oxyradicals, binds zinc-finger repair proteins (39) and inhibits nucleotide excision repair (55). Separately, AQP3 transports hydrogen peroxide (21) and supports peroxide-dependent signaling in lung adenocarcinoma(67), a route needing no arsenite, and AQP3 gating responds to pH and peroxide itself (57), making the two coupled rather than independent. Our data limit the second: AQP3 alone raised basal damage only as a trend and N-acetylcysteine removed nothing without arsenite. One transcriptional observation favors the first: SQSTM1, whose product sequesters KEAP1 and thereby activates NRF2, is the only gene in the ten-cell design significantly raised above both single conditions where a genuinely combined oxidant insult should appear.

Several limitations qualify the study. Intracellular arsenic was never measured, so the influx inference rests on the mutant series and published transport phenotypes (24, 25, 27, 68). Construct expression was never quantified against endogenous AQP3, no loss-of-function arm was run, and no damage-endpoint arm above 5 μM was acquired. γH2AX reports a single S/G2-biased response (34–36) rather than double-strand breaks specifically, both responsive lines are virally transformed, and the transcriptional design is plate-confounded, which leaves that interaction untested rather than rejected. One caveat runs the other way: aquaglyceroporins are bidirectional, and AQP9-null mice clear arsenic poorly and are more rather than less sensitive to arsenite (20), so "more aquaglyceroporin, more damage" is an in vitro statement.

Nor does anything here speak directly to human risk: 1 μM arsenite is about 75 ppb, roughly seven and a half times the United States maximum contaminant level (7), applied to transfected, sorted, SV40-immortalized fibroblasts for 72 h. Against those limits, the design does what it set out to do. The interaction rests on a within-date paired design in which each construct is compared only with the control beside it; the cell-type dependence is reproduced with three paralogues in one assay and host-cell set; and the effect follows the AQP3 sequence rather than the act of expressing a membrane protein. What this system provides is not an estimate of human risk but a modifier that can be switched on and off while the exposure-response relationship is re-measured in the same cells, the manipulation that human association studies (12, 14) cannot perform. Further work will add ICP-MS across the mutant series, loss-of-function and expression controls, and per-cell γH2AX read against per-cell AQP3 level.

## DATA AVAILABILITY

SHERRY and Duplex-seq sequencing data will be deposited and made available in the NCBI Sequence Read Archive (SRA) upon publication. All other data supporting the findings of this study are included in the article and its supplementary Information. Code used in this study is available from the corresponding author upon reasonable request.

## ACKNOWLEDGEMENTS

We thank the Creighton University Flow Cytometry Core for flow cytometry support and Texas A&M University High Performance Research Computing for computing support.

## CRediT authorship contribution statement

**Shiwei Yin:** Investigation, Methodology, Validation, Formal analysis, Visualization, Writing – review & editing. **Bingru Feng:** Software, Formal analysis, Data curation, Visualization, Writing – review & editing. **Maryam Vaziripour:** Investigation, Methodology, Validation, Writing – review & editing. **Shannon E. Slewitzke:** Investigation, Validation, Writing – review & editing. **Gail F. Fernandes:** Investigation, Validation, Writing – review & editing. **Jihye Yun:** Resources, Methodology, Writing – review & editing. **Christopher I. Amos:** Conceptualization, Resources, Writing – review & editing. **Jun Xia:** Conceptualization, Methodology, Formal analysis, Data curation, Visualization, Supervision, Project administration, Funding acquisition, Writing – original draft, Writing – review & editing.

## SUPPLEMENTARY FIGURE LEGENDS

**Supplementary Figure S1.**
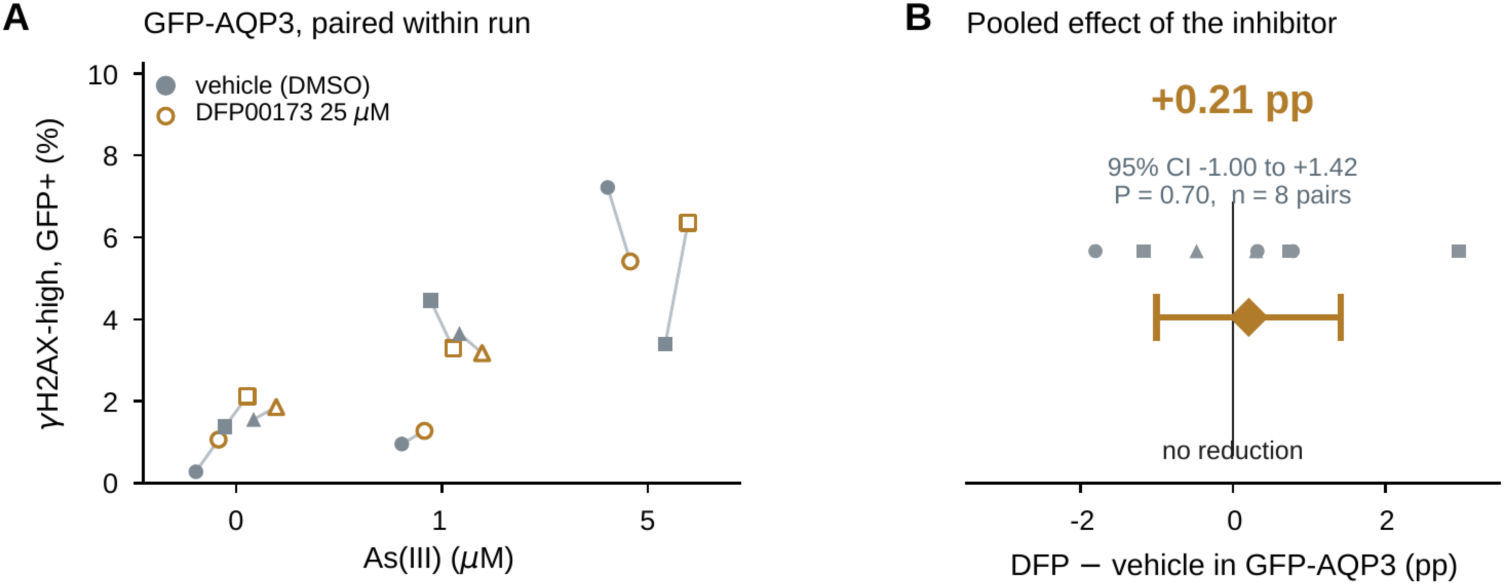
DFP00173 does not reduce the AQP3-associated signal. DFP00173 is an AQP3-selective inhibitor. If the γH2AX excess in AQP3-expressing cells required flux through the channel, blocking the channel should lower it. Only GFP-AQP3 is shown: the GFP-Tubulin arm of this experiment transfected at 3.8–11.7% GFP+ (against 20–23% on the dose-series plates) and its untransfected cells already carried 1.3–11.4% γH2AX-high background, so it reports well condition rather than construct and is omitted. The contrast is within construct and within run: one GFP-AQP3 transfection split between DMSO vehicle and 25 μM DFP00173, read on the same plate at the same arsenite concentration. (A) Every vehicle/DFP pair, joined; symbol shape identifies the acquisition run. (B) The pooled paired difference over all eight pairs: +0.21 pp, 95% CI −1.00 to +1.42 pp, two-sided paired t-test P = 0.70. Diamond, the mean; bar, the 95% CI; grey symbols above, the eight individual pairs. Pairing within run removes the run-to-run variation that dominates the raw values in A, which is why this interval is narrow on eight observations. Conclusion. DFP00173 at 25 μM does not reduce the γH2AX signal in AQP3-expressing MRC5-SV40 cells. The pooled change is +0.21 pp and the 95% interval excludes any reduction larger than 1.0 pp. That bound is smaller than the AQP3-over-Tubulin excess at 1 μM (2.58 pp) and far smaller than at 5 μM (7.69 pp, Figure 8G), so removal of the channel-attributable signal is excluded at those doses; at 0 μM, where the excess is 1.51 pp, the interval reaches the boundary and full removal is only marginally excluded. Unchanged signal under a channel inhibitor is consistent with the mutant data (Figure 3), where the pore mutant H53F lowers the signal and the phosphorylation-site mutant S152A does not. Tests here are two-sided, as for the other control analyses in this paper. n = 8 paired observations across three acquisition runs.

**Supplementary Figure S2.**
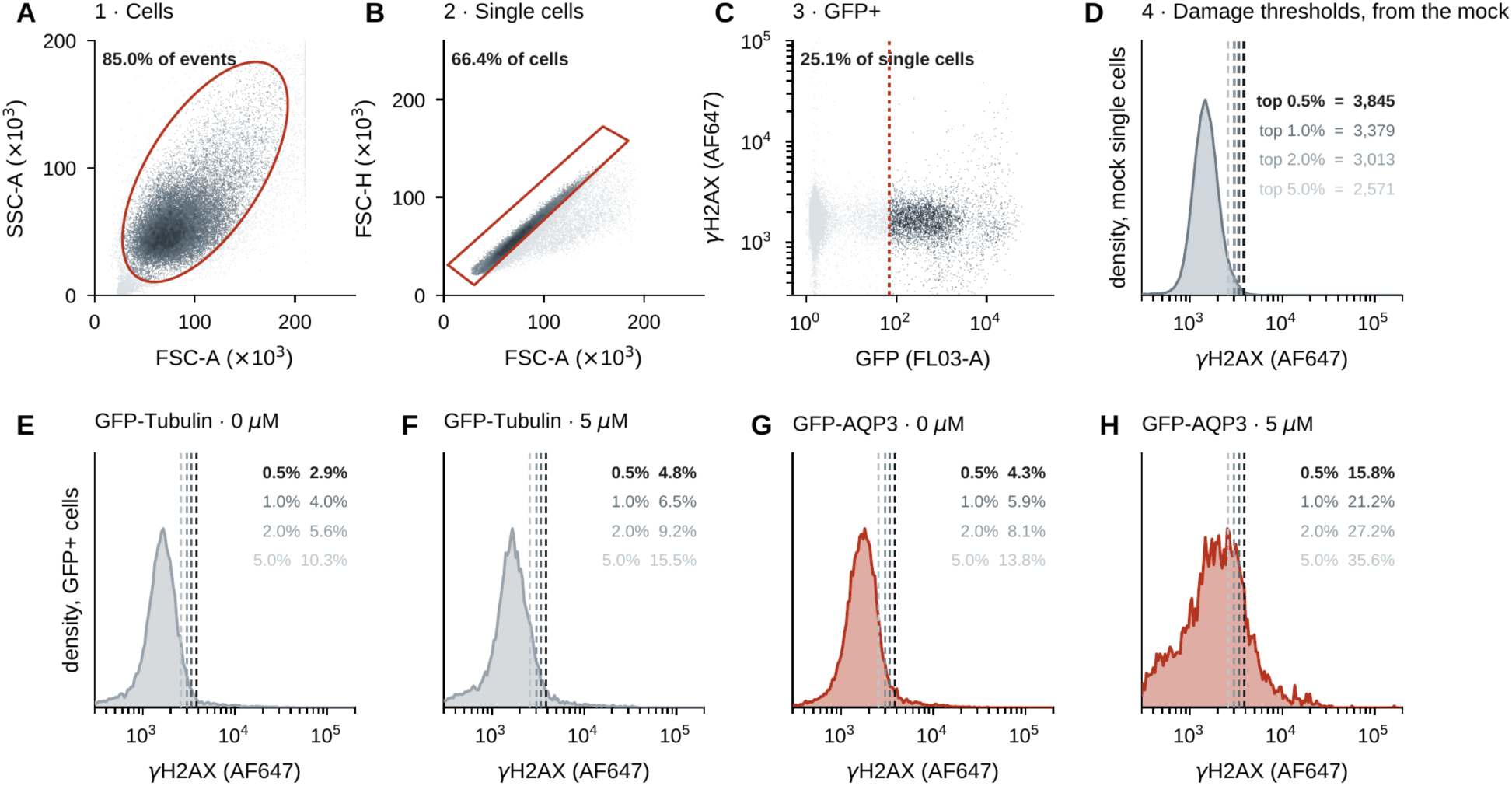
Gating strategy, gate by gate, and the mock-anchored thresholds. MRC5-SV40, 15 September 2023. The lab’s FlowJo gates were read out of the workspace file and applied event by event in Python; the outlines drawn here are those stored gates, not redrawn approximations, and the percentages are of the parent population, as FlowJo reports them. (A) Cells gate on FSC-A × SSC-A (85.0% of recorded events). (B) Single cells on FSC-A × FSC-H within the cells gate (66.4% of cells). (C) The GFP gate within single cells (25.1% of single cells), shown against γH2AX. Grey, events outside the gate; shaded dots, events retained; red outline or dotted line, the gate. (D) Where the damage thresholds come from: the mock-transfected, antibody-stained single-cell distribution of that acquisition, cut at its top 0.5, 1, 2 and 5% (3845, 3379, 3013 and 2571 in AF647 units on this plate). A mock-anchored threshold re-derived for every acquisition absorbs day-to-day differences in staining and instrument gain, which a fixed cutoff would not. (E–H) All four cutoffs applied to the GFP-positive cells of four representative tubes, with the percentage above each. The 0.5% cutoff is the pre-specified primary; every contrast in the paper is reported at all four, and the threshold sweeps in Figures 1, 4 and 5 show that no conclusion depends on which is used. In panels A–C each event is drawn as one dot, shaded by local dot density.

**Supplementary Figure S3.**
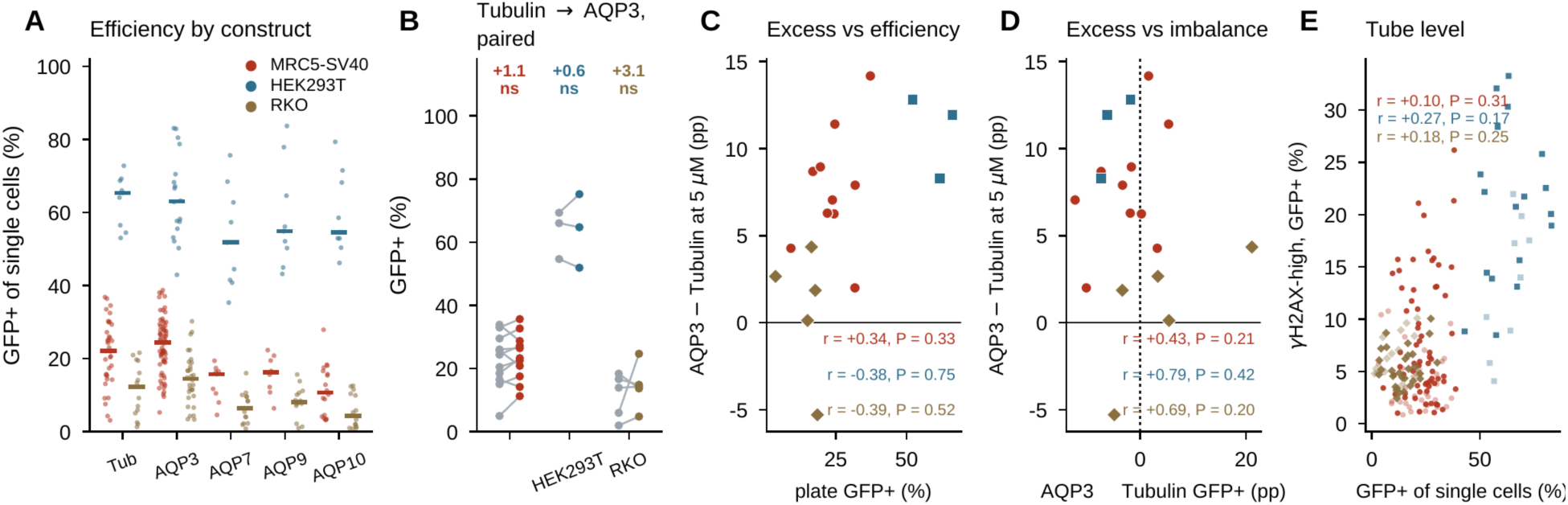
The damage phenotype is not a transfection-efficiency artifact. Three arguments, none of which relies on the others. (A) GFP-positive fraction of single cells for every gated tube of the arsenite γH2AX experiments, by construct within each line; bars, medians. Efficiency differs a great deal between lines (median 65% in HEK293T, 22% in MRC5-SV40, 12% in RKO) and between constructs. (B) The two arms that are actually compared are matched. Within an acquisition plate, GFP-AQP3 exceeds GFP-Tubulin by +1.1 pp in MRC5-SV40 (n = 11 plates, P = 0.34), +0.6 pp in HEK293T (n = 3, P = 0.84) and +3.1 pp in RKO (n = 5, P = 0.50); none is distinguishable from zero. (C, D) The size of the AQP3-over-Tubulin excess at 5 μM does not track how well the plate transfected (r = +0.34, −0.38, −0.39; P = 0.33, 0.75, 0.52) nor how unbalanced the two arms were on that plate (r = +0.43, +0.79, +0.69; P = 0.21, 0.42, 0.20). None is significant and the signs do not agree across lines. (E) At tube level the readout is not a function of efficiency: pooling GFP-Tubulin and GFP-AQP3 within a line, r = +0.10, +0.27 and +0.18 (P = 0.31, 0.17, 0.25).

**Supplementary Figure S4.**
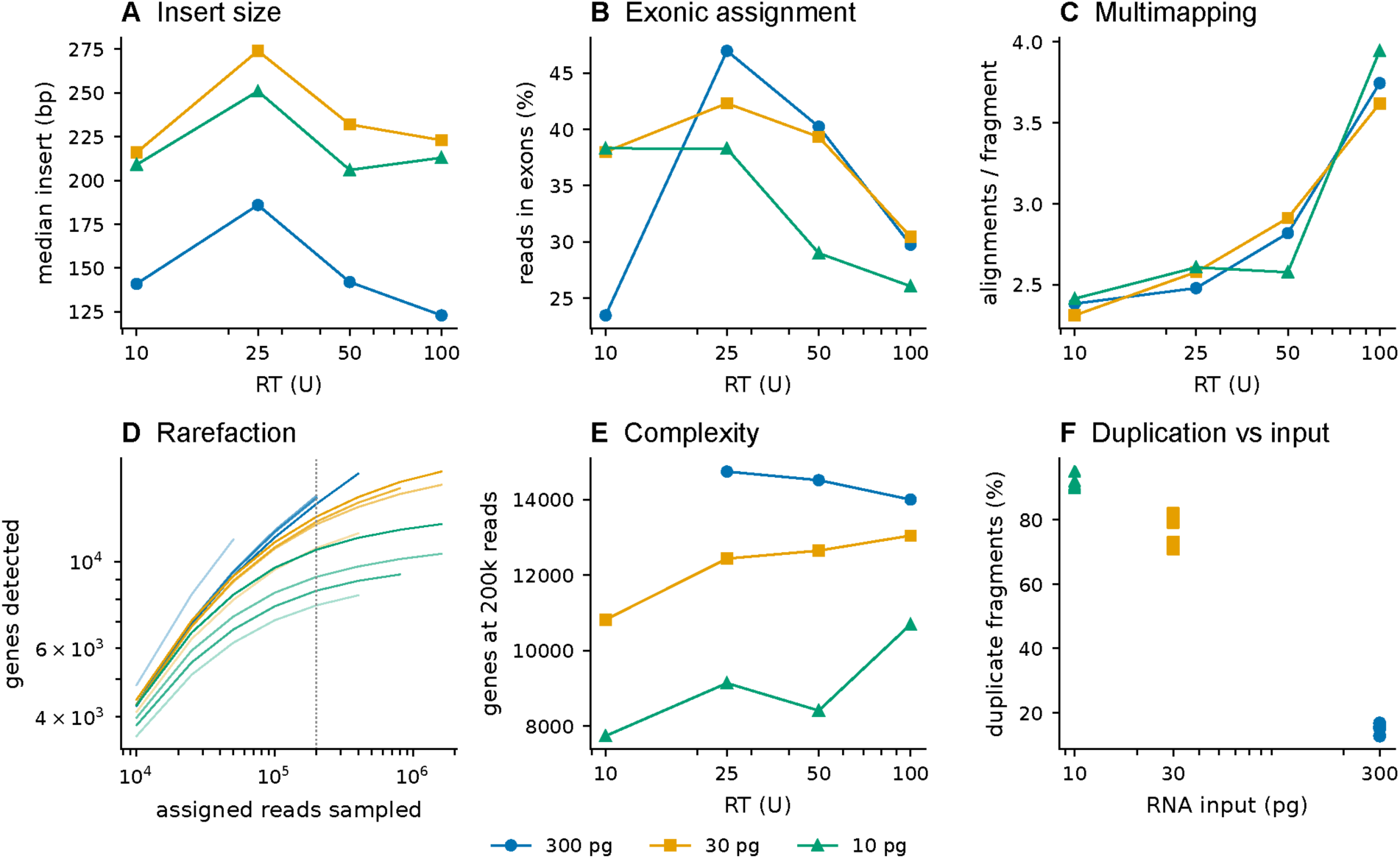
SHERRY optimization in HEK293T cells. A complete 3 × 4 factorial of RNA input (300, 30 and 10 pg) against reverse transcriptase (10, 25, 50 and 100 U), one library per condition, with Tn5 held constant; colour gives RNA input. (A) Median insert size. (B) Exonic assignment rate, reads assigned to exons as a share of counted reads. (C) Alignments per fragment, a measure of multimapping. (D) Rarefaction: genes detected against assigned reads sampled without replacement, opacity increasing with reverse transcriptase; dotted line, the 200,000-read depth used in E. (E) Genes detected at a matched depth of 200,000 assigned reads; the 300 pg, 10 U library is absent because it was sequenced below that depth. (F) Duplicate fragments against RNA input. Above 25 U the added enzyme buys off-target product rather than transcriptome: alignments per fragment rise and exonic assignment falls at 100 U in all three input rows, and insert size peaks at 25 U although Tn5 is constant, so the extra material is short, spuriously primed product. Sensitivity and specificity trade off against input: at 300 pg, 25 U is best on every metric (14,742 genes at matched depth against 14,000 at 100 U), whereas at 30 and 10 pg depth-matched complexity is still rising at the top of the range (13,046 and 10,692 genes at 100 U), so conversion rather than specificity limits those libraries and 100 U was used for the single-cell and ten-cell preparations. Duplicate fraction separates the three inputs without overlap (13– 17%, 71–82% and 90–95% at 300, 30 and 10 pg), which confirms the input labels independently of the read data. One library per condition, so there is no within-condition replication and no error bar: every statement is consistency of direction across the three input rows rather than a test. Three further libraries prepared at 100 U by a different operator were excluded, for short inserts (102–107 bp), low exonic rate (18.9–29.6%) and a duplicate fraction that rose rather than fell with RNA input.

